# Endophytic and ectomycorrhizal, an overlooked dual ecological niche? Insights from natural environments and *Russula* species

**DOI:** 10.1101/2024.01.24.576884

**Authors:** Liam Laurent-Webb, Philippe Rech, Amélia Bourceret, Chloé Chaumeton, Aurélie Deveau, Laurent Genola, Mélanie Januario, Rémi Petrolli, Marc-André Selosse

## Abstract

- Ectomycorrhizal (EcM) fungi play key roles in ecosystem functioning, in particular temperate ones. Recent findings suggest that they can endophytically colonize the roots of non-EcM plants. Here we aim at (i) providing new evidence of colonization of non-EcM hosts by EcM fungi, (ii) exploring factors driving such colonization (plant identity, site, root filter), and (iii) providing direct microscopical evidence for endophytism.
- Using amplicon sequencing (ITS2), we described the root fungal communities of 42 plant species collected at nine locations in France. In two of those sites, we also compared rhizosphere and root fungal communities to identify a potential root filter. Finally, we investigated endophytism in *Russula* spp. at two *Russula*-rich sites using fluorescence *in situ* hybridization (FISH) paired with confocal microscopy.
- We find a large but variable share of EcM sequences in roots of non-EcM plant species, in particular nearby EcM hosts, suggesting that endophytism is a secondary ecological niche. Though EcM fungi were more abundant in the rhizosphere compared to roots, their composition was similar to that of roots, suggesting a poor root filter. We observed metabolically active hyphae of *Russula* spp. endophytically colonizing the apoplast of two non-EcM plant species.
- As shown for other EcM fungi (*e*.*g*., *Tuber* spp., Ascomycota) we demonstrate the dual EcM/endophyte niche for *Russula* (Basidiomycota). The ecological consequences of this duality still need to be addressed. The ability to colonize two ecological niches may be a trait kept by EcM fungi which evolved from endophytic fungi, as stipulated by the “waiting room hypothesis”.

## Introduction

Most plant species form root mutualistic associations with soil fungi, the mycorrhizas (van der Heijden *et al*., 2015), where water and nutrient foraged in soil by fungi are exchanged for plant photosynthates (Smith and Read, 2009). Ectomycorrhizal (EcM) fungi form a hyper-diverse, polyphyletic group of great importance especially for trees in temperate forest ecosystems (Tedersoo & Brundrett, 2017). Beyond mycorrhizal fungi, roots are also loosely colonized by various biotrophic fungi, which, unlike mycorrhizas, do not form a well-structured and elaborated interface of exchanges. These so-called endophytes range from latent pathogens to true mutualists (Wilson, 1995; Rodriguez *et al*., 2009).

There is scattered, but increasing evidence that some EcM fungi colonize endophytically the roots of non-ECM plant species (*i*.*e*., which are either non-mycorrhizal or mycorrhizal with arbuscular mycorrhizal fungi). For instance, EcM Sebacinaceae species are common plant endophytes (Selosse *et al*., 2009; Weiß *et al*., 2016), EcM *Tricholoma matsutake* endophytically colonize roots of non-EcM plants *in vitro* (Murata *et al*., 2013, 2014), and EcM *Inocybe* spp. can abound in roots of nettle (*Urtica dioica*) growing under poplar trees (Yung *et al*., 2021). Barcoding, gene expression and FISH microscopical analyses support endophytic growth of truffles in non-EcM plants (*Tuber* spp.; Schneider-Maunoury *et al*., 2018; 2020), perhaps with slightly deleterious effects on these host (Taschen *et al*., 2020). Finally, many EcM species are phylogenetically intermingled with endophytic species, *e*.*g*. in the Trechisporales (Vanegas-León *et al*., 2019), Helotiales (Wang *et al*., 2006), Pyronemataceae (Hansen *et al*., 2013; Hughes *et al*., 2020), Hygrophoraceae (Halbwachs *et al*., 2018) or Hymenochaetales (Korotkin *et al*., 2018). Here, we aim at (i) providing new evidence of colonization of non-EcM host by EcM fungi, (ii) exploring factors driving the colonization of non-EcM plants by EcM fungi (plant identity, site, root filter), and (iii) providing direct microscopical evidences for endophytism.

### Detection of EcM fungi in roots of non-EcM plants

We harvested healthy roots from non-EcM plants in France at four locations distant from 20 to 450 km (three forest sites and one meadow) and five neighboring sites distant from 10 to 800 m (two forest sites, one close meadow and two intermediate edge sites, near Gaillac city; Fig. S1 and Methods S1). From these nine sites, we barcoded the root mycobiota of 433 non-EcM plant individuals belonging to 42 species (557 root samples; Fig. S2; Table S1; see molecular methods in Methods S2). We detected molecular barcodes of EcM fungi in roots of non-EcM plants (Fig. 1) in 69% of investigated plant individuals, in which > 1% of reads belonged to EcM fungi. They accounted for 2.1 to 35% of the reads in forest/edges sites *versus* 0.37 to 1.1% in meadows, suggesting that the presence of EcM plant host favored the settlement or the competitivity of EcM fungi in adjacent non-EcM hosts. EcM fungal community composition varied significantly between sites (Table S3; see Methods S3 for statistical methods), even at meter scale (*e*.*g*., MF1 and F4 sites separated by 10 meters; Fig. 1b). Such spatial turnover was often reported for EcM fungi colonizing EcM hosts (*e*.*g*., Richard et al., 2005). In particular, Fabaceae and Cyperaceae were abundantly colonized in edges and forest sites but not in meadows (Fig. S3b), suggesting again that the presence of EcM hosts enhances their colonization.

**Figure 1:**
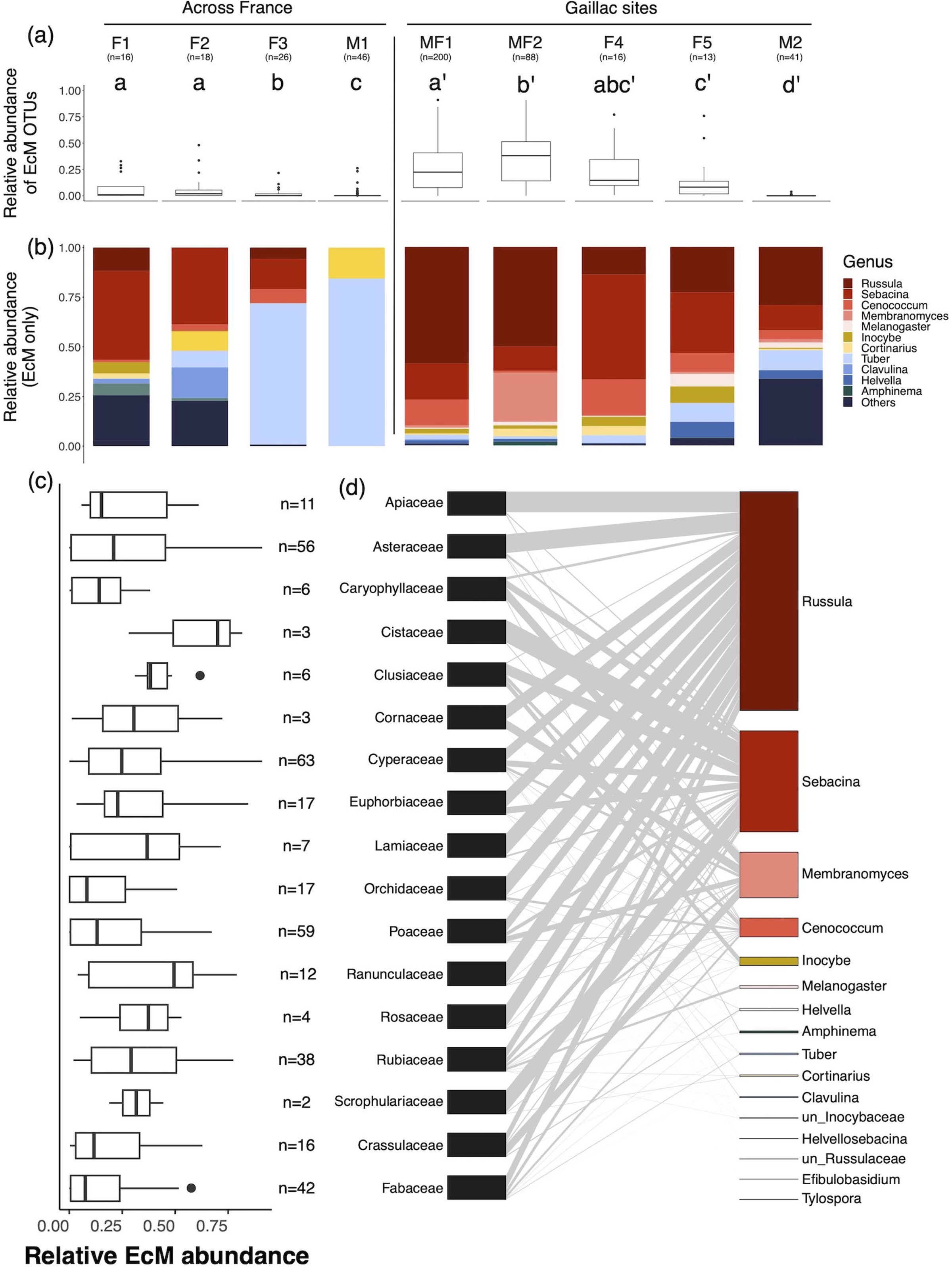
Ectomycorrhizal fungi are found as endophytes in roots of non-EcM plants across sites and plant families. **(a)** Relative abundance (proportion of reads) of EcM fungi in the total fungal community. Different letters indicate significant differences (pairwise Wilcoxson test, *p*<0.05) between sites across France or between Gaillac sites, separately. Tests were performed separately for sites across France and Gaillac sites. **(b)** Composition of the EcM community in roots of non-EcM plants at the genus level in the 9 sites. **(c)** Relative abundance of EcM fungi per plant family at Gaillac sites. The number of samples considered for each family is indicated at the right of each boxplot. **(d)** Bipartite network of interactions between plant families and EcM genera at Gaillac sites.

The abundance of EcM fungi varied between families, with no clear taxonomic signal, and none of them was completely devoid of EcM fungi (Fig. 1c; Fig. S3). EcM community composition displayed significant differences between plant families: depending on the statistical method used, 12%-13% of the variability was explained by plant families in Gaillac sites, and 34-36% in other sites across France (Table S2). Therefore, the ability to host EcM fungi may not be equally distributed among plants, deserving further study. We noted that the two forest edges (MF1 and MF2) sites in Gaillac displayed a high abundance of *Russula* spp. (Fig. S4), respectively accounting for 59 and 50 % of EcM reads in these sites, and contrasting with the forest sites where *Sebacina* are overrepresented. This parallels the fact that the forest edges sites displayed abundant *Russula maculata* and *R. delica* fruitbodies (our personal observations).

The bipartite networks of interactions between non-EcM plants and EcM fungi (either in Gaillac sites, or across France, Fig. 1c and Fig. S5 respectively) harbor similar characteristics in comparison to available knowledge on true EcM interactions (van der Heijden *et al*., 2015). Indeed, the Gaillac network (local scale) was moderately modular (Q=0.33) and specialized (H_2_’=0.53; Table S3; see Methods S3.3 for null models) compared to the network of sites across France (regional scale) which was highly modular (Q=0.59) and specialized (H_2_’=0.81). Patterns of nestedness were contrasted: the network of sites across France was significantly anti-nested while the Gaillac one was significantly nested (Table S3). Nestedness in bipartite networks analysis is thought to be characteristic of mutualistic interactions (Bascompte *et al*., 2003), but recent approaches showed that it is a poor predictor of mutualism (Pichon et al., 2023; see Discussion S1). Thus, EcM fungi are ubiquist in non-EcM plants but display high levels of network modularity and specialization, two correlated network properties (Dormann & Strauss, 2014) which may result from one-sided or reciprocal adaptation between EcM fungi and non-EcM plants (Dormann et al., 2017; see Discussion S1).

### Non-EcM roots poorly filter EcM fungi

In two forest sites (F1 and F2), rhizospheric soils paired with previously described non-EcM roots of seven species were additionally collected and barcoded (Fig. S6a). EcM reads were 3 to 135 times more abundant in rhizosphere than in the roots (Fig. 2a), suggesting that rhizosphere is a more propitious niche than roots for EcM fungi. The percentage of EcM OTUs simultaneously found in both roots and rhizosphere varied across plant species (26%-65% of OTUs, which represented 66%-99% of the total EcM reads Table S4). On the contrary, rhizospheric EcM OTUs undetected in roots (25-73% of OTUs) were lower in abundance (<1 to 34% of EcM reads; Table S4). The EcM diversity in roots and rhizosphere was not significantly different (Fig. S6b). While the EcM fungal community composition did not differ significantly between roots and rhizosphere (Fig. 2c; Table S6), the whole community of fungi, whatever their ecology, significantly differed between the two compartments (*PERMANOVA*; *p*<10^−4^ Fig. 2b; Table S6), as often reported due to root filtering (Gottel *et al*., 2011; Maciá-Vicente *et al*., 2020; Bourceret *et al*., 2022). The filtering of EcM fungi is thus lower than that of the total fungal community, which is a strong biological signal that non-EcM plants poorly filter EcM fungi.

**Figure 2:**
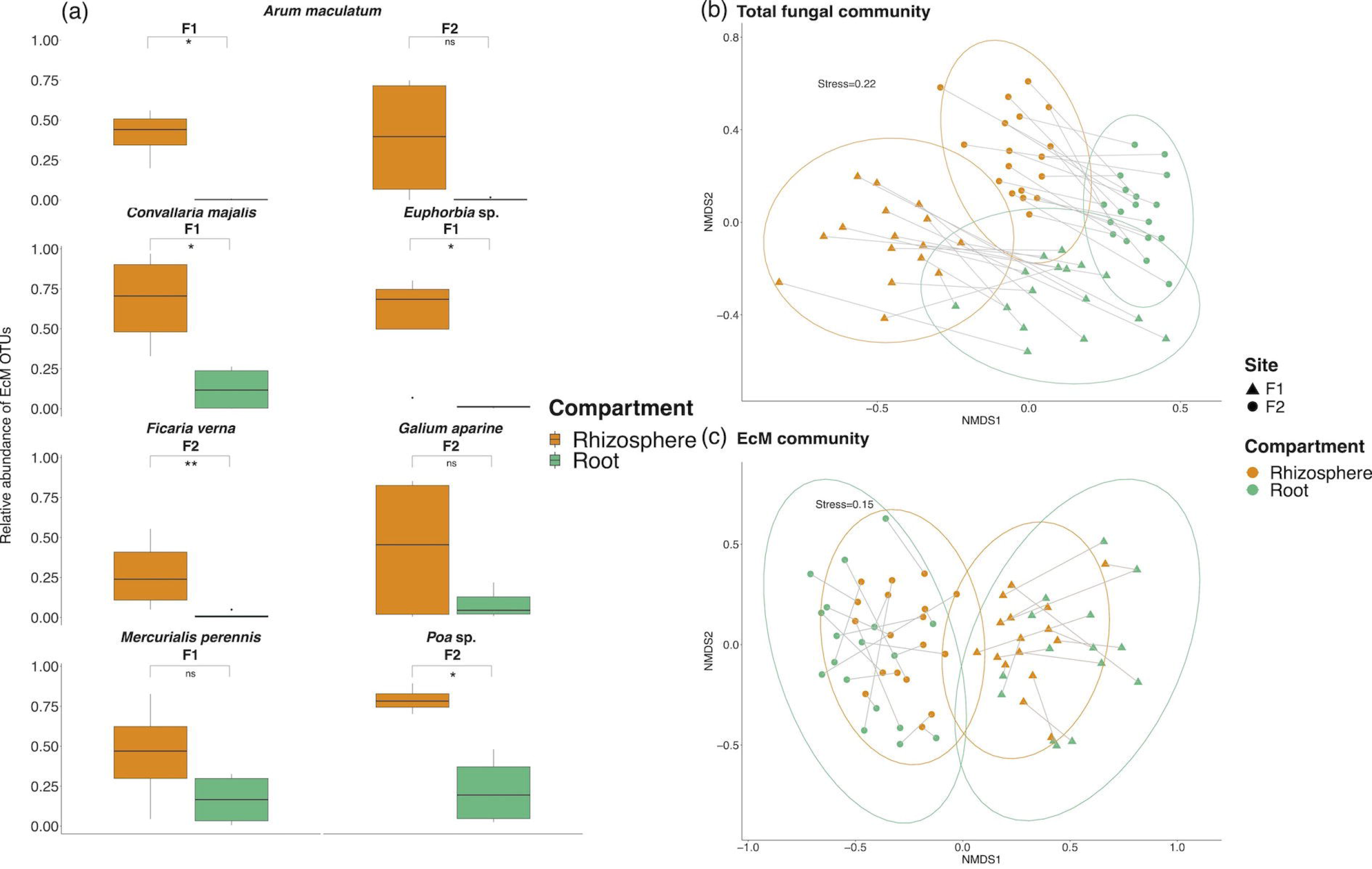
The EcM fungi colonizing roots of non-EcM plants are scarce, but of similar composition compared to rhizospheric soil. **(a)** Boxplot of the total relative abundance in rhizosphere (orange) and roots (green) of EcM fungi by plant host species in the sites F1 and F2. Significant differences between rhizosphere and roots were tested using a Wilcoxson test. n.s: p>0.05; *: p<0.05, **: p<0.00. **(b), (c)** NMDS of the Bray-Curtis distance matrix (based on relative abundances) for the total fungal community **(b)** and the EcM community only **(c)**. Circles represent the normal distribution of the data. NMDS stresses are indicated on the top left of each NMDS. Pairs of samples are connected by grey lines.

Though total root and rhizosphere EcM communities are not significantly different, some EcM genera are significantly differentially abundant in the two compartments (Fig. S7). *Cenococcum, Lactarius* and unidentified Cortinariaceae are more abundant in roots while some genera are more abundant in the rhizosphere (*e*.*g*., *Inocybe, Clavulina*; Fig. S7). However, the main EcM genera in these sites did not display significant differences between roots and rhizosphere (*e*.*g*., *Sebacina*). Whether the differences between EcM fungi are linked to their niche preferences between compartments and/or to root filtering deserves further studies. Differences in colonization success observed between EcM fungi may be linked to species-specific genomic features, in particular Plant Cell Wall Degrading Enzymes (PCWDE) as observed for endophytes (Mesny *et al*., 2021). Though *Inocybe* spp. are significantly more abundant in rhizospheres compared to roots in our study, some authors found that they colonize non-EcM species (Yung *et al*., 2021).

Overall, this low filtering may explain the limited differences in EcM fungal community observed between rhizosphere and roots in the plant species from our study. Though we observed a low root-filtering of rhizospheric EcM fungi, EcM root communities are mainly explained by host identity: this apparent discrepancy suggests that host filtering already occurs in both rhizosphere and roots (Discussion S2).

### Direct evidence for endophytism in *Russula* spp

Barcoding is sensitive to contamination, although the various signals reported above advocates against general contamination. In addition, EcM fungi rarely produce asexual spores, and thus scarcely contaminate samples. Yet, the reads of EcM fungi reported above in roots may result from surface colonization of roots rather than internal, true endophytism. We took profit of sites MF1 and MF2 abundantly colonized by *Russula* spp., a common EcM genus (Hackel *et al*., 2022), to directly visualize and confirm the root-mycelial interactions *in situ*, as in Schneider-Maunoury *et al*. (2020). We investigated the presence of *Russula* spp. in non-EcM roots by fluorescence *in situ* hybridization (FISH), paired with confocal microscopy. We designed two probes specifically targeting *Russula* spp. 18S rRNA based on sequences publicly available and sequences from fruitbodies harvested in the Gaillac sites MF1 and MF2 (Material S4.1). *In silico* and *in vitro* controls confirmed that (i) the probe RUS899 hybridizes with *Russula* spp. (Table S6) but also with the closely related EcM *Lactarius* spp., while (ii) the probe RUS101 is specific for *Russula* spp. although it does not hybridize with all species (Fig. S8-9). As no *Lactarius* sequences occurred in Gaillac barcoding results, both probes are suitable to visualize *Russula* spp. In 32 samples from 5 non-EcM plant species (Table S7) we revealed all hyphae based on cell wall coloration by SR2000 and the generalist fungal probe EUK516 (Fig. 3a; Material S4), as well as *Russula* spp. hyphae with our specific probes. We observed *Russula* spp. hyphae in the roots of two species, *Carduus pycnocephalus* (Asteraceae; Fig. 3b-d; Fig. S10) and *Ranunculus bulbosus* (Ranunculaceae; Fig. S11), which both are arbuscular mycorrhizal (Mullen and Schmidt, 1993; Torrecillas *et al*., 2012). Hyphae are observed on the root surface, intimately anchored to the epidermis cell walls, and some are clearly situated within the apoplast of root cortical cells (*e*.*g*., Fig. 3b-d). No signs of necrosis, parasitism, nor apparent tissue differentiation were observed in investigated roots. The colonization by *Russula* spp. is thus endophytic (*sensu* Wilson, 1995). The observed hyphae are alive and metabolically active since we labelled their rDNA, but we also observed inactive hyphae (*i*.*e*., with cell wall stained but without cytoplasmic labelling; Fig. 3a).

**Figure 3:**
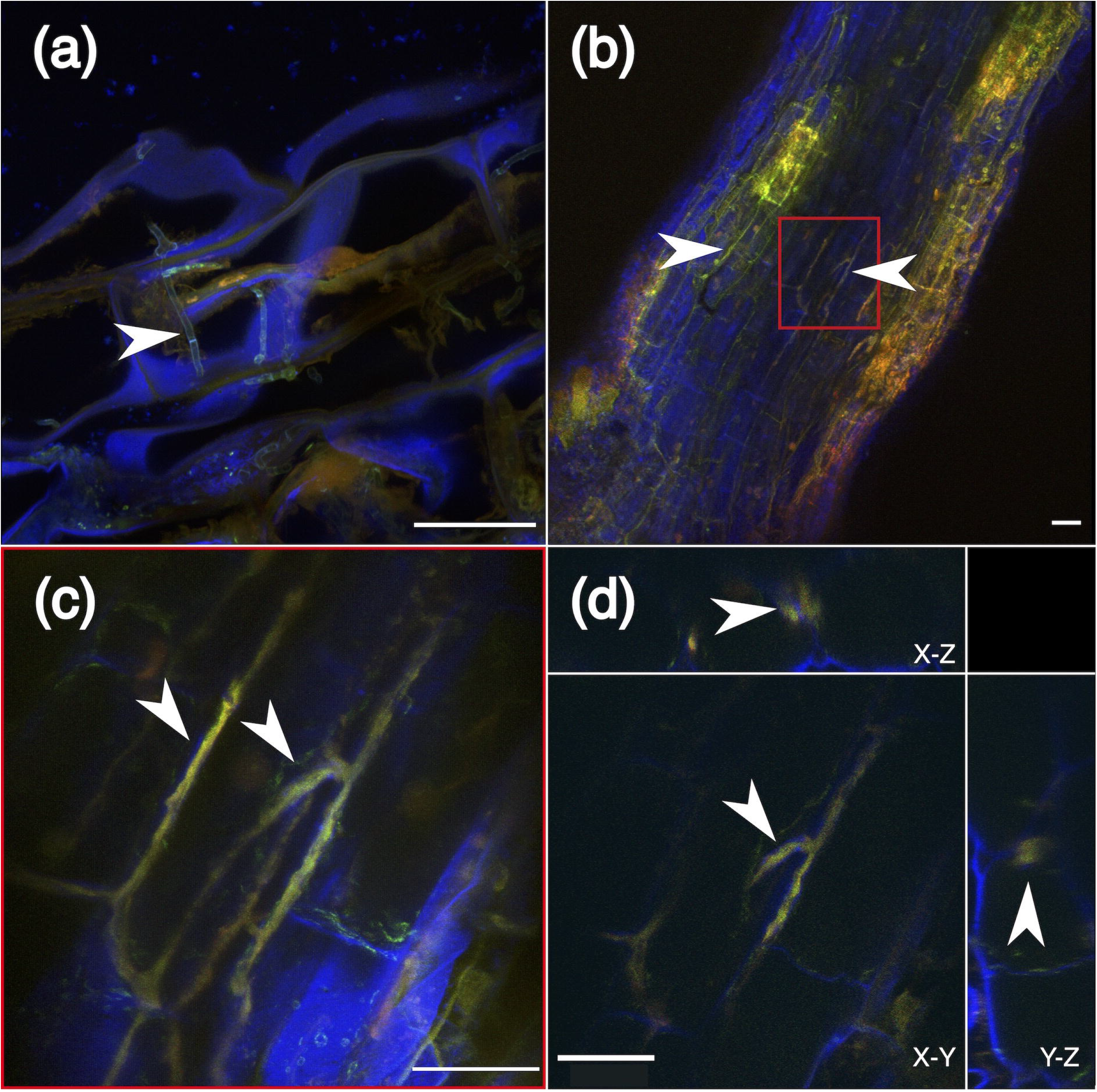
Hyphae of *Russula* spp. colonize roots of non-EcM plant species. **(a)** Fungal hyphae stained with the cell wall marker SR2000 (blue signal) in a negative control (no probes). Green and red signals correspond to autofluorescence produced mainly by cell walls of fungi (white arrow) and plant. **(b)** Fungal hyphae only hybridized with the generalist probe (EUK516-ATTO565) (left arrow) and *Russula* spp. hyphae hybridized with the generalist and *Russula* specific (RUS101-ATTO633) probes (co-hybridization appears in orange) in roots of *Carduus pycnocephalus*. **(c)** Red area from (b) magnified (63x) to show *Russula* spp. hyphae inside the roots of a *C. pycnocephalus* individual collected in MF1 site (Gaillac). Hyphae colonize the apoplast of plant cells. Maximum intensity projection of a z-stack. **(d)** Orthogonal representation of the same area. Scale bars: 30 μm.

### Conclusion and perspectives

Our results further support that EcM fungi can colonize roots of non-EcM plants, especially in the vicinity of EcM trees. Bipartite networks analyses reveal high levels of modularity and specialization, which may arise from one-sided adaptation or co-evolution processes (Dormann *et al*., 2017). Finally, we visually confirm endophytism in non-EcM roots for *Russula* species.

The physiological role of EcM fungi in roots of non-EcM plants is yet to be determined, especially whether they are parasitic, commensal or mutualistic, given that hyphae are metabolically active (Schneider-Maunoury *et al*., 2018; 2020 for *Tuber;* this work for *Russula*). Taschen *et al*. (2020) showed in mesocosms that the development of non-EcM plants was lower when colonized by *Tuber melanosporum*, either due to parasitism of non-EcM plants, or to a competition for root colonization between EcM fungi and the root mutualists (*e*.*g*., arbuscular mycorrhizal fungi). Deleterious effects of EcM fungi on non-EcM plants partially explain patches of reduced vegetation observed around trees producing EcM fruitbodies of *Tuber* (Splivallo *et al*., 2012), which was also observed in this study for the *Russula*-rich sites (our personal observations at MF1 and MF1 2; Fig. S1c). Different EcM species may differ in their effects on plants, if any. From a fungal point of view, non-EcM roots could provide resources to EcM fungi. Indeed mycelium development of *T. melanosporum* is favored by non-EcM plants (Taschen *et al*., 2020). Yet, the low abundance of EcM fungi in meadow sites suggest that this ecological niche is hardly sufficient, and at most adds up to the resources from EcM roots. One may even speculate that their abundance at edge sites (MF1 and 2) may be one player in the replacement of non-EcM plants by EcM trees in ecological successions.

The colonization of non-EcM plants by EcM fungi illustrates an example of ecological plasticity increasingly acknowledged in fungi (Selosse *et al*., 2018). Such plasticity could be inherited from the evolution processes called the ‘waiting-room hypothesis’ (van der Heijden *et al*., 2015; Selosse *et al*., 2021) which stipulates that the ectomycorrhizal association evolved from endophytism. Indeed, some fungi may have retained endophytic abilities while developing complex mutualistic interactions such as ectomycorrhizas, leading to such dual endophytic + EcM ecological niche. Be it vestigial or not, this endophytism deserves further studies addressing its ecological importance as well as its potential to shape functional networks created between EcM and non-EcM plants.

## Acknowledgements

M-A.S. acknowledges the Institut Universitaire de France for financial support. Molecular work was performed at the BoEM laboratory (MNHN). The authors thank Laure Schneider-Maunoury and Bastien Bennetot for their help with samples collection and molecular processing.

## Authors contribution

M-A.S. designed and supervised the work; L.L.-W. performed the experimental work, analyzed the data and drafted the manuscript with M-A.S.; A.B. supervised the molecular work with L.L.-W.; P.R., C.C. and A.D. supervised the microscopy work that also involved M.J. for acquisition and L.L-W. for acquisition and images analysis; R.P. helped in experimental work and analysis and L.G. provided access to the Gaillac sites; all authors corrected the final manuscript.

## Data availability

18S sequences of *Russula* spp. fruitbodies harvested *in situ* are available at NCBI (https://www.ncbi.nlm.nih.gov/) under accession numbers OR910623-OR910639. *Metabarcoding* data are also available at NCBI under SRA project number PRJNA1067284.

## Competing interests

No competing interests.

## Notes

### Competing Interest Statement

The authors have declared no competing interest.

