## Supplemental Information for "Endophytic and ectomycorrhizal, an overlooked dual ecological niche? Insights from natural environments and *Russula* species"

### Supplementary Methods (1-4)

#### **<u>Supplementary Methods 1:</u> Study sites and sampling**

This study encompasses results from nine sampling sites presented below (Fig. S1; Table S1).

##### **<u>Methods S1.1:</u>** ***Endophytism of EcM fungi in non-EcM plant species in several sites across France***

We sampled non-EcM plant individuals in three forest sites (F1, F2 and F3) and one meadow (M1) across France in order to test whether the dual endophyte/EcM niche is a common feature observed in sites separated by 20 to 450km (Fig. S1). These samples were collected between May 2016 and April 2019 in four locations (Table S1). We sampled roots of non-EcM plants in two forests characterized by an acidic soil dominated by chestnut trees (*Castanea sativa*; F1 and F2 sites). At both sites, the four most abundant herbaceous species were selected and several individuals were sampled (Table S1). Four root tips per individual were pooled. We additionally collected rhizospheric soil in order to characterize a potential root filter on rhizosphere EcM fungi. Rhizospheric soil of each individual was collected by gently shaking the soil closely attached to roots. Root samples of the F3 site were collected on a calcareous field in Corrèze, near but outside a productive *Tuber melanosporum* orchard. M1 samples were collected in a meadow in a closed botanical garden of the National Museum of Natural History in Paris and 4 root tips per non-EcM plant individual were collected and treated individually (i.e., no pooling). All collected root tips (F1, F2, F3 and M1 sites) were selected based on their apparent good health and checked for the absence of ectomycorrhizae (especially root tips of ligneous species). They were surface sterilized as follows: root tips were washed three time in 70% ethanol (3x5min) and in 0.9% bleach. They were then rinsed three times in sterile water and stored at -20°C before molecular analysis.

##### **<u>Methods S1.2:</u>** ***Detection of EcM fungi in roots of non-EcM plants by amplicon sequencing and fluorescence* in situ *hybridization in Gaillac sites***

In order to investigate the variability of the dual EcM/endophyte niche at smaller scale, we sampled roots from non-EcM plants species in 5 sites near Gaillac in South-West of France, distant from 10 to 800m. The sites are characterized by a calcareous soil. We collected samples in (i) two forests dominated by oaks, where fruitbodies of *Russula* spp. and other EcM fungi have been observed (F4 and F5), (ii) a meadow where no fruitbodies were observed (M2), and (iii) areas at the edge of the forests with only non-EcM plant species but producing a large number *Russula* spp. fruitbodies (MF1 and MF2; Fig. S1; Table S1).

Samples were collected at different times: in June 2021, we collected roots only for amplicon sequencing (Methods S2) in the five sites (MF1, MF2, F3, F5 and M2). The sampling was representative of the plant community within each site, and therefore varied between them. We sampled individuals of the same species in the different sites whenever possible. For each plant individual, three root tips were collected, carefully washed in sterile Mili-Q water, pooled and kept dried in silica-gel. Samples were kept dry until molecular work. We also collected *Russula* spp. fruitbodies in the forest edge site MF1 for molecular identification and fluorescence *in situ* hybridization probe design (see Methods S4.1 for probe design). In September 2021, we collected plant individuals for both amplicon sequencing and fluorescent *in situ* hybridization (*FISH*) microscopy in both edges sites (MF1 and MF2). We carefully dug up individuals in order to keep the roots in their surrounding soil. Samples were kept at 4°C in individual zip-locks. The next day in the laboratory, we removed the rhizospheric soil and selected three root tips per individual. Each root tip was cut in two pieces: one piece was carefully washed in sterile water, flash-frozen in nitrogen and kept at -20°C before molecular analysis. The other half was washed in PBS 1% and fixed in PFA 4% to fix RNAs overnight at 4°C before *FISH* and microscopy observations (Methods S4). Fixed samples were kept at -20°C before hybridization. In September 2021, one *Russula* sp. fruitbodie was also collected and processed for *FISH* in order to test our newly designed probes (see Methods S4). Finally, we sampled individuals of two plant species (*Ranunculus bulbosus* and *Pilosella officinarum*) for *FISH* in July 2023 and processed them as described above.

#### <u>Supplementary Methods 2:</u> Sequencing of fruitbodies and amplicon sequencing of root/rhizospheric fungal communities

##### <u>Methods S2.1:</u> *Fruitbodies gDNA extraction and 18S/ITS2 rRNA PCR*

We sequenced the 18S and ITS region of the ribosomal operon of *Russula* spp. fruitbodies harvested in Gaillac sites to design fluorescence *in situ* hybridization probes (18S) and for taxonomic identification (ITS). 20 to 80 mg of dried (in silica-gel) fruitbodies were used to extract gDNA. Samples were mechanically disrupted 3 times at 30 hz in a TissueLyser II (Qiagen, USA) during 30 sec with 2 grinding steel balls (Retsch, Luxembourg). gDNA was extracted using the commercial DNeasy Plant Mini kit (Qiagen, USA) according to the manufacturer's instructions, and eluted in 100µl of TE buffer. Extracted gDNA was quantified using Qubit™ fluorometer (Life Technologies, Singapore) and Qubit™ (1X dsDNA High Sensitivity Assay Kit, Invitrogen). 18S rDNA was amplified from each fungal gDNA extraction by PCR using the universal fungal primers NS1 (3'-GTAGTCATATGCTTGTCTC-5') and NS8 (3'- TCCGCAGGTTCACCTACGGA-5'; White *et al.*, 1990). The ITS1-5.8S-ITS2 rDNA region was amplified using primers ITS5 (3'-GGAAGTAAAAGTCGTAACAAGG-5') and ITS4 (3'-TCCTCCGCTTATTGATATGC-5'; White *et al.*, 1990). PCR reaction was performed by mixing 3ng of gDNA, 7.5pmole of each primer, 2U DFS-Taq DNA Polymerase (Bioron Life Science, Germany), 75µg Bovine Serum Albumin (Sigma, USA), 200µM of each dNTP (New England Biolabs), and 1x Incomplete NH4 (Bioron Life Science, Germany). The optimal conditions for PCR amplification of 18S rDNA segments with these primers were 95°C 10min, followed by 30 cycles at 95°C 30 sec, 53°C 30 sec, 72°C 2 min, and a final elongation step at 72°C 10 min. The PCR products were subjected to Sanger sequencing from both directions using the same set of primers (Eurofins Genomics, Germany). Additionally, a couple of internal primers were designed specifically on conserved regions with the free software Primer3 (Untergasser *et al.*, 2012) to sequence the extremities of each 18S PCR fragments. The obtained electropherograms were checked, sequences were corrected and submitted to GenBank under the accession number OR910623-OR910639. ITS sequences taxonomy was assigned against the UNITE online database using BLAST tool (https://unite.ut.ee/analysis.php#).

##### <u>Methods S2.2:</u> *Amplicon sequencing of the fungal communities*

Root and rhizosphere samples from the F1, F2, F3 and M1 sites were processed as in Schneider-Maunoury *et al.* (2018, 2020). Briefly, the ITS2 region of the ribosomal operon was amplified using the primers ITS86-F (3'-GTGAATCATCGAATCTTTGAA-5') and ITS4 (see above) which favor the detection of Ascomycota and Basidiomycota (the two phyla containing ectomycorrhizal fungi) over Glomeromycota (Waud *et al.*, 2014; Op De Beeck *et al.*, 2014). PCR products were purified, pooled in equimolar pools and sequenced with an Ion Torrent sequencer (Life Technologies, Carlsbad, USA).
Samples from Gaillac sites were disrupted with a TissueLyser II using two inox beads (3x30s at 30 Hz). Total gDNA was then extracted using the Qiagen Plant Kit following the manufacturer instructions and

stored at -20°C. We also performed extractions with no root samples as negative controls. All gDNA extracts were purified to remove potential PCR inhibitors. To do so, gDNA was mixed with Ampure XP (Beckman Coulter Inc., USA) solution in a 1 (gDNA) : 1.8 (Ampure) (v:v) ratio. The reaction plate was then placed on a magnetic plate to separate beads from the solution. The supernatant was removed and beads were washed with 70% EtOH two times. gDNA was then resuspended in EB buffer. Purified gDNA concentrations were measured with PicoGreen and all gDNAs were diluted to 3.5 ng/µL. The ITS2 region of the risobomal operon was amplified using barcoded ITS286-F/ITS4 primers in triplicate with the DFS-Taq (Bioron Life Science, Germany). The PCR mixture consisted in 2.5 µL of Incomplete Buffer (10x), 0.5 µL of $MgCl_2$, 2.5 µL of 3% BSA, 0.5 µL dNTPs (10 mM each), 0.75 µL of each primer (at 10 µM), 2U of Bioron DFS-Taq, 3 µL of DNA (3.5 ng/µL) and 14.1 µL of $H_2O$. The cycling conditions were as follow: 10 min at 95°C (initial denaturation), 35 steps of denaturation (95°C, 30s), annealing (56.5°C, 30s), elongation (72°C, 30s) and a final elongation at 72°C for 10 min. We also added PCR negative controls for which DNA was replaced by water. PCRs products were then checked on a 2% agarose gel and purified using AMpure XP solution with a 1:1 (v:v) ratio as described before. Purified PCRs products concentration was measured with PicoGreen and pool in equimolar quantity. Equimolar pool was also purified. Sample pools were sequenced using a 2x250 Miseq technology on an Illumina platform by Fasteris SA (Switzerland).

**<u>Methods S2.2:</u>** ***Bioinformatic analysis of amplicon sequences***

As fungal communities of samples from sites across France (F1, F2, F3 and M1) and Gaillac sites (MF1, MF2, F4, F5, M2) were obtained using two different sequencing technologies, the first processing steps were slightly different. Samples from the first four sites were sequenced in distinct IonTorrent runs. Reads were then assembled and demultiplexed on the IonTorrent platform. Sequences were quality checked, with less stringent parameters in order to keep a large sequencing depth (*--fastq_maxns* 1, *--fastq_maxee* 3, *--fastq_minlen* 200). For samples from Gaillac sites, paired-end reads were assembled and quality checked (*--fastq_maxns* 0, *--fastq_maxee* 2) using cutadapt (Martin, 2011) and then demultiplexed (keeping only sequences longer than 200 pb).
Assembled and demultiplexed reads of the 9 sites were then processed altogether. Reads were dereplicated and clustered as classical 97% sequence similarity Operational Taxonomic Units (OTUs) using VSEARCH (Rognes *et al.*, 2016). Sequences were checked for the presence of chimeras (*--uchime_denovo*). The taxonomy was assigned with VSEARCH against the Unite v8.3 database (Nilsson *et al.*, 2019). Reads were filtered in order to keep only non-chimeric sequences of > 200 pb, with a total abundance of at least 10 and a spread (i.e., the number of samples where the read is present) superior or equal to 1. We used the *decontam* algorithm (Rivera *et al.*, 2011) to remove the potential contaminants in the Gaillac samples, using both algorithms (*prevalence* and *frequency*). First, we used the *prevalence* algorithm using the negative extraction and PCR controls with a stringent threshold of 0.5. We then used the *frequency* algorithm using default parameters. A few more filters were applied to all datasets:

we removed samples less than 2 000 fungal reads, OTUs with less than 5 reads per sample and OTUs representing less than 0.5% of the reads per sample.

We inferred functional fungal traits using the FUNGuild database (Nguyen *et al.*, 2016). We only considered “Probable” and “Highly Probable” assignations and classified the others as “Unknown”. OTUs assigned to multiple trophic guilds (e.g., Saprotroph-Plant Pathogen) were grouped in the category “Others”.

#### <u>Supplementary Methods 3:</u> Statistical analysis of fungal communities

Amplicon sequencing data were processed using the R software (R Core Team, 2023) and the package *phyloseq* (McMurdie and Holmes, 2013).

**<u>Methods S3.1:</u>** ***Influence of the sites and plant host on fungal communities***

In order to test for differences in EcM relative abundance between sites, we used pairwise Wilcoxson rank test (*pairwse_wilcoxson_test*, rstatix package; Kassambara, 2023). We used β-diversity metrics to determine the influence of the location and the plant host family on total and EcM fungal communities. To do so, we computed Bray-Curtis distances from both relative abundances, as they may perform better for community comparisons (Gloor *et al.*, 2017; McKnight *et al.*, 2019), and Hellinger-transformed data to correct for variability in sampling depth (Legendre & Gallagher, 2001). We computed PERMANOVA with 10,000 permutations using the *adonis2* function of the *vegan* R package (Oksanen *et al.*, 2013). The formula used was: *distance matrix ~ site* x *family*. Analyses were run separately for the four sites across France and Gaillac sites as fungal communities were sequenced with different technologies.

**<u>Methods S3.2:</u>** ***Differential abundances analyses of OTUs between environment types in Gaillac sites***

In order to test whether some fungal OTUs were significantly more abundant in one of the three environments of the Gaillac sites (*i.e.*, forest, meadow and forest edge), we ran several differential abundances analysis as advised in Nearing *et al.* (2022): *LEfSE* (Segata *et al.*, 2011), *ANCOM-BC* (Lin & Peddada, 2020) and *ALDEx* (Fernandes *et al.*, 2013). OTUs were considered differentially abundant in one environment only if they were identified as differentially abundant with the three methods. We represented differentially abundant OTUs based on the Linear Discriminant Analysis score (LDA) computed in the *LEfSE* procedure.

**<u>Methods S3.3:</u>** ***Constructing and characterizing bipartite networks of interactions between EcM fungi and non-EcM plants***

We built bipartite networks in order to search for patterns of associations between EcM fungi and non-EcM-plants and to compare their structure to available knowledge on true mycorrhizal associations. We used the *bipartite* R package (Dormann *et al.*, 2008) to build plant/EcM fungi bipartite networks. We built the networks at both fungal genera/plant family and fungal OTUs/plant species levels to see if observed patterns were consistent across different taxonomic levels. We built one network for the four sites across France (regional scale) and one for Gaillac sites (local scale). Furthermore, we built one network for each site separately to ensure that observed patterns are not linked to differences between sites, in particular modularity and specialization which can be linked to differences between habitats. We plotted the bipartite graphs using the *plotweb* function (*bipartite* package). To characterize the networks' structure, we computed several metrics: (i) nestedness (weighted NODF, *networklevel* function), (ii) modularity (Q, Beckett's algorithm from the *computeModules* function) and (iii) specialization ($H_2$'; *H2fun* function). (i) In nested networks, specialists interact mainly with generalists and *vice versa* (Bascompte *et al.*, 2003). (ii) Modularity arises from subsets of preferential associations, which may be linked to reciprocal adaptations within modules (Dormann *et al.*, 2017). (iii) Finally, $H_2$' measures the level of specialization in interactions among plant and fungus and allows comparisons between networks of different size (Blüthgen *et al.*, 2006). $H_2$' varies between 0 (no network specialization) to 1 (network fully specialized). In order to test whether the properties of the bipartite networks were significantly different than expected by chance (that is, different from random patterns of associations), we generated, for each network, 1,000 random networks using the *quasiswap* null model algorithm (*permatswap* function), a stringent algorithm that keeps both connectance and marginal sums of the original network constant. Networks are considered nested (or anti-nested) if more than 97.5% of the generated networks have a lower (resp. higher) weighted NODF than the real network ($p<0.025$). Similarly, networks are considered significantly modular and specialized if more than 97.5% of the generated networks have lower Q and $H_2$' values, respectively ($p<0.025$).

**<u>Methods S3.4:</u>** ***Comparisons of rhizosphere and root EcM fungal communities***

In order to quantify and characterize a potential root filter applied to rhizospheric EcM fungi, we compared the EcM $\alpha$-diversity between compartments for each species by computing the Shannon index and tested for significant differences between compartments using Tukey's Honest Significant Differences after running the following linear regression: *Shannon index ~ plant species* x *compartment*. Normality and homoscedasticity of the Shannon index were checked using diagnostic plots. We also compared the composition of the total and EcM fungal communities in the two compartments using PERMANOVA (*adonis2* function of the *vegan* R package, 10,000 permutations) with the following model: *distance matrix ~ compartment* x *plant species*. As detailed before, distance matrices were computed using the Bray-Curtis distance based on both relative abundances and Hellinger-transformed data. Differences in composition were represented using Non-Metric Multidimensional Scaling (NMDS). Pairs of samples are connected by a grey line. For each species, we identified the number of OTUs shared (or not) between the two compartments and computed their relative read abundance in the EcM community (threshold of one read). Finally, we assessed if some EcM fungal genera are significantly more abundant (or not) in one of the two compartments using differentially abundant analysis. To do so, we used the Linear Discriminant Analysis Effect Size procedure (*LEFsE*; Segata *et al.*, 2011).

#### **Supplementary Methods 4: Fluorescence *in situ* hybridization (*FISH*) and microscopy detection of *Russula* hyphae**

**Methods S4.1:** ***Design of rRNA probes specific to* Russula *spp. for* FISH *and microscopy observations***

Oligonucleotide probes were designed to target the 18S rRNA of *Russula* genus including the *Russula* fruitbodies harvested in areas producing *Russula* capropohores in the Gaillac sites. *Russula* 18S ribosomal DNA (rDNA) sequences were obtained from fruitbodies harvested on and outside productive areas and from genomic data published in NCBI GenBank or Mycocosm databases (The Fungal Genomics Resource-JGI) (cf. Supp. File 1 to see accessions used). *Russula* 18S rRNA sequences were aligned using the free Multalin algorithm (Corpet, 1988) along with 18S rRNA sequences of a selection of several orders in Agaromycetes and Russulaceae family. The alignment was scanned visually to detect regions of sequence homology suitable for *Russula* genus-specific probes, i.e., conserved within the *Russula* genus but with differences (mismatches) compared to other genera. We identified 6 regions exhibiting polymorphisms which could therefore be used as a basis for designing probes targeting *Russula* spp. (or including closely related genera in the Russulaceae family). Physical accessibility of probes to these regions was assessed using reference maps of probe accessibility and sensitivity available in *Prymenisum parvum* (Metfies and Medlin, 2008). The specificity of potential probes was tested *in silico* using iterative BLAST searches with low stringency algorithm parameters in the GenBank and SILVA databases. Taking into account the number of mismatches of the designed probes and the *in silico* accessibility and specificity, we selected 2 probes: RUS899 and RUS101 (see table below for sequences). *FISH*-probes were commercially synthesized by Biomers (Biomers.net, Ulm/Donau, Germany) including 5′-end labeled with ATTO fluorochromes and stored in sterile DNA-grade water at -20°C.

| **Name** | **Sequence** | **Fluorescent dye** |
|---|---|---|
| EUK516 | ACCAGACTTGCCCTCC | ATTO565 |
| NON-EUK516 | GGAGGGCAAGTCTGGT | ATTO565 |
| RUS101 | ATGTAGAAAGGTATCATCAAAT | ATTO633 |
| RUS899 | CGCAATAGTTTGTCTTGCGTAAAT | ATTO633 |
| NON-RUS899 | ATTTACGCAAGACAAACTATTGCG | ATTO633 |

**Methods S4.2:** **In vitro** ***specificity test of the newly designed probes***

Strains of *Phanerochaete chrysosporium* RP78, *Lactarius quietus* S05C, *Russula luteotacta* 201811 RL, *Russula* sp. AT 0111 were cultivated on P5 medium (Paschelwski agar medium 5, Di Battista *et al.*, 1996) in order to test probes' sensitivity and specificity *in vitro* and to set up confocal acquisition parameters. All strains were supplied by the UMR IAM 1136 isolates collection grown for several days in P5 medium at 25°C. When fungal elements became visible by eye, the hyphae were harvested, washed with PBS 1x, fixed in the 3% PFA fungi fixation solution for 24 h (see Supp. File 2 for details on all reagents used), washed in PBS and finally stored at -20°C in 50% ethanol-PBS 1x until use. Moreover, different part of fruitbodies harvested *in situ* (cape and/or stipe) of *Russula* spp. and two other undetermined fungi species (C1 and C2) harvested on and outside forest edges were included in the specificity test. *Russula* sp. (stipe and cape), and undetermined fungi C1 and C2 were razor blade cut, rinsed in PBS 1x and fixed in the fungi fixation solution as for pure culture fungi. For *FISH*, fruitbodies and mycelium samples were gradually dehydrated in an ethanol series prepared in sterile ultrapure water (50%, 70%, 96% and 100%) 5 min each and rehydrated in an ethanol series (70%, 50%, 30%, 10%) prepared in PBS-T 5 min each before a last PBS-T bath. Samples were treated for 15 min at 30°C with FCWE digestion solution to weaken fungal cell walls before prehybridization at 46°C during 10 min. The prehybridization buffer was then replaced with a hybridization buffer containing the probe(s) (0.35pmol/µL of hybridization solution each) during 2h at 46°C. After a stringent wash in the washing solution during 10 min at 46°C followed by PBS 1x bath, sample were mounted in antifade solution, and visualized on a laser scanning confocal microscope (Zeiss LSM980; Carl Zeiss, Oberkochen, Germany) equipped with an Airyscan2 detector and coupled to ZEN Blue 3.3 software (Carl Zeiss). In some cases, the cultivated hyphae have been placed and fixed on Poly-L-lysine coated slides (Sigma-Aldrich, St. Louis, MO, USA) before applying the *FISH* protocol. Fungal samples were co-hybridized with either the universal eukaryote probe EUK516 and with one of the following probes: sense *Russula* probes (RUS899/RUS101), non-sense *Russula* probe (NonRUS899) or none sense EUK516 probe, using a combination of distinct fluorescent dyes.

**<u>Methods S4.3:</u>** ***Hybridization and observation of* Russula *spp. hyphae in roots of non-EcM plant species***

To visualize *Russula* fungi within non-EcM plant roots, we set up a *FISH* experiment. Roots samples harvested in September 2021 and July 2023 (Table S1) were washed carefully several times in water and PBS 1x in order to eliminate the rhizospheric soil. Root pieces collected were cut into two consecutive segments of 1-cm long. One was immediately immersed in the plant fixation solution (PFA 4%) and incubated overnight at 4°C to fix RNAs. Samples were then rinsed three times in PBS 1x and progressively dehydrated in a series of ethanol solutions (10%, 30% and 50%) during 15 min each. The root segments were stored in the sample storage solution and kept at -20°C for further analyses. Segments for which *Russula* sequences were detected (Methods S2) were used for *FISH* experiments. We used two types of probes, both targeting 18S rRNA. One is the universal eukaryote probe EUK516, which targets the 18s rRNA of eukaryotic cells (Amann *et al.*, 1990; see above for probes sequences) coupled with ATTO-565 dye (Biomers, Germany), hereafter called “EUK565”. The other probe was either one of the two newly designed probes targeting specifically *Russula* 18S rRNA (RUS889 and RUS101, Methods S4.1) coupled with ATTO-633 dye (Biomers, Germany). As in Schneider-Maunoury *et al.* (2020), fluorochromes were chosen to minimize the autofluorescence of the fungi and plant root tissues. In order to visualize plant and fungal cell walls, the dye SR2000 (Renaissance Chemicals) was added to the hybridization buffer.

Selected samples were treated for 1h at 30°C with PCWE digestion cocktail 1x mixed with FCWE digestion solution 1x to weaken plant and fungal cell walls. After rinsing them into PBS-T, samples were then treated with 0.08µg/µl of Proteinase K during 30min at 37°C. The proteinase K reaction was stopped by replacing the solution by Glycine working buffer during 2 min and the samples were incubated 30 min in the post fixative solution, rinsed and prehybridized at 46°C during 30 min. The prehybridization buffer was replaced with hybridization buffer containing the probe(s) (0.35pmol/µL of hybridization solution each) during 2h at 46°C. After a stringent wash in the washing solution during 10 min at 46°C, sample were mounted in antifade solution mixed with 0.1% of SR2000 (Renaissance Chemicals).

The *FISH*-stained root samples were further mounted with Slowfade Diamond Antifadent (Molecular Probes, Eugene, OR, USA, Thermofisher, cat. no. S36963) and stored at 4°C over night until observation with a laser scanning confocal microscope (LSM 980 Zeiss microscope) equipped with an EC-Plan Neofluar 10x/0.3 dry objective, a W Plan-Apochromat 20x/1.0 water and a Plan-Apochromat 63x/1.4 oil objective. The Airyscan 2 module was used for acquisitions using the multiplex 4Y mode. Probes labelled with ATTO565 or ATTO633 dyes were excited at 568nm or 647 nm wavelengths respectively and detected with a bandpass filter (570-630 nm and 660-720 nm). The cell wall fluorescent dye SR2000 was visualized with a laser excitation at 405 nm wavelength and recorded with a bandpass filter (420-477nm). For each field of view, an appropriate number of optical sections were acquired with a Z-step of 0.15 to 1 µm. Airyscan images were reconstructed and analysed using ZEN Blue 3.5, ZEN 2.1 LITE black software (Zeiss), the Vision4D 3.0.1 software (arivis AG, Germany) or the free software FIJI-ImageJ. Hybridization signals obtained on non-ECM roots samples were successfully and specifically detected for EUK516 and the sense *Russula* probes and the same parameters (laser power and gain of detector) were used to image the control samples: samples without any probe or samples hybridized with the non-sense probes.

### Supplementary discussions (1-2)

**<u>Supplementary discussion 1:</u> The structure of mutualist interactions' bipartite networks**

In order to further characterize the interactions between EcM fungi and non-EcM plants, we constructed the network of interactions between EcM fungi and non-EcM plants in Gaillac sites (local scale; Fig. 1d), in sites across France (regional scale; Fig. S5), and in each site individually. We assessed the topological properties of the networks by computing the commonly used modularity and nestedness (here weighted nestedness; Bascompte *et al.*, 2003; Olesen *et al.*, 2007). Modularity (that is, groups of species that interact preferentially) may arise in intimate interactions such as symbiotic mutualism or parasitism, resulting in reciprocal specialization (Olesen *et al.*, 2007) but may also be linked to environmental or geographic factors. Nestedness was thought to be a characteristic of mutualistic networks (Bascompte *et al.*, 2003) providing stability in the community. In a nested network, generalists tend to associate with specialists and *vice versa*. However, recent results show that nestedness is not reliable to predict if a network is mutualist (Pichon *et al.*, 2023 and citations hereafter).

Here, we show that the colonization of EcM fungi in roots of non-EcM plants results in a modular network in both Gaillac sites (Fig. 1d; Table S3) and sites across France (Fig. S5; Table S3). At the site level, modularity was frequently observed especially in networks at the fungal OTUs/plant species level (Table S3). Patterns of nestedness, non- and anti-nestedness were observed across sites with no clear tendency. The network of Gaillac sites was significantly nested (Table S3) while the network of sites across France was significantly anti-nested. Different mycorrhizal types tend to have contrasted network structures. For instance, arbuscular mycorrhizal fungi and plants tend to form nested networks (Chagnon *et al.*, 2012; Montesinos-Navarro *et al.*, 2012) with intermediate modularity (Toju *et al.*, 2014; Põlme *et al.*, 2018). EcM interactions on the contrary are often reported to form non- or anti-nested networks (Bahram *et al.*, 2014; Toju *et al.*, 2014; Roy-Bolduc *et al.*, 2016) though nestedness was observed in some cases (Peay *et al.*, 2007). Modularity was sometimes observed in EcM/plant networks but is not systematic (see previous studies). Anti-nestedness and modularity were also observed in epiphytic orchids mycorrhizal interaction networks (Martos *et al.*, 2012; Petrolli *et al.*, 2022) and in ericaceous plant fungus networks (Toju *et al.*, 2016). Thus, mutualistic mycorrhizal associations result in diverse networks topologies depending on the mycorrhizal type(van der Heijden *et al.*, 2015). In this work, we find that EcM fungi colonizing non-EcM plants result in modular bipartite networks, with contrasted patterns of nestedness, fitting observations in true EcM fungi.

Furthermore, we assessed the specialization of EcM fungi and non-EcM plants networks using the $H_2$', allowing comparisons between networks of different sizes (Blüthgen *et al.*, 2006). We found high degrees of specialization in both Gaillac sites and sites across France (Table S3). As with modularity, specialization was often observed at the individual site level, in particular at the fungal OTUs/plant species level. Observed specialization was greater than in some mycorrhizal associations (Toju *et al.*,

2014; Perez-Lamarque *et al.*, 2022) but similar to that observed in grasses root endophytes (Kivlin *et al.*, 2022).

In this study, bipartite networks of interactions between non-EcM plant roots by EcM fungi are both modular and specialized, two correlated network properties (Dormann & Strauss, 2014). These properties were also observed within sites suggesting that they are not only linked to differences between sites but rather to preferential interactions between plants and fungi (Dormann *et al.*, 2017). These patterns may arise from several process such as one-sided adaptation or co-evolution (Dormann *et al.*, 2017).

#### <u>Supplementary discussion 2:</u> A rhizospheric host filtering of EcM fungi by non-EcM plants?

We assessed the EcM community of roots of non-EcM plants from several families in different sites. Comparisons of EcM communities show significant differences between plant families (Fig. 1c; Fig. S3; Supp Table 2c,d). Furthermore, bipartite networks of interactions between non-EcM plants and EcM fungi are highly specialized and modular (Fig. 1d; Fig. S5; Table S3), even at the site level (Table S3), suggesting preferential interactions between groups of EcM fungi and non-EcM plants. Altogether, these results suggest a host filtering of EcM fungi. However, the comparison of rhizosphere and roots EcM communities in paired samples revealed no significant differences in EcM community composition between the two compartments (Fig. 2; Fig. S6; Table S5), though EcM fungi were much less abundant in roots than in rhizosphere (Fig. 2a).

This apparent discrepancy suggests that host filtering of EcM communities by non-EcM plants may occur both in rhizosphere and in roots. First, root exudates may shape EcM community composition in the rhizosphere. Knowledge acquired in true EcM associations demonstrate that plant exudates such as flavonoids or abietic acid may participate to EcM development and spores' germination in the rhizosphere (as reviewed in Garcia *et al.*, 2015; Martin *et al.*, 2016). In this study, we investigated roots (and rhizosphere) of plants forming arbuscular mycorrhizae (AM) or no mycorrhizae (NM) for which molecular dialogue with EcM fungi has not been investigated. As EcM plants, AM plants produce root exudates such as strigolactones that participate to AM fungal partners' recruitment (Bonfante & Genre, 2010), and thus shape the rhizosphere microbiome (Uroz *et al.*, 2019). It has been hypothesized that strigolactones may also be involved in EcM molecular dialogue (Garcia *et al.*, 2015) despite conclusive evidence. The rhizosphere is a complex and dynamic compartment within which root and fungal exudates are likely to influence the development of fungi (*e.g.*, EcM fungi) and thus influence root mycobiota assembly, leading to differences between plant species.

Second, the ability of EcM fungi to colonize non-EcM roots may be limited, leading to a low abundance of EcM fungi in non-EcM roots compared to the rhizosphere (Fig. 2a). The fungal colonization in true mycorrhizal associations usually requires an important root remodeling to ensure access of root tissues to the fungal partner and reduced plant defenses. Reduction of plant defense during colonization is modulated by fungal exudates (*e.g.*, Small Secreted Proteins in EcM associations; Plett *et al.*, 2014; Martin *et al.*, 2016) ensuring a specific recognition of mycorrhizal fungi. The endophytic niche may facilitate and predispose plant and fungal adaptations allowing more specialized and complex interactions such as mycorrhizae, as stipulated by the 'waiting-room' hypothesis (Selosse *et al.*, 2009, 2021).

### Supplementary Tables (1-7)

**<u>Supplementary Table 1</u>: Details on samples collected.**

Location, number of root samples, number of individuals, number of plant species/family and their corresponding analysis.

| Location | Name | Date | Nb. root samples | Nb. individuals | Nb. species | Nb. family | Rhizospheric soil | Analysis |
|---|---|---|---|---|---|---|---|---|
| Orry-la-Ville (ORY) | Forest 1 (**F1**) | April 2019 | 18 | 18 | 4 | 3 | 18 | Ion Torrent sequencing (ITS2) |
| Orsay (ORS) | Forest 2 (**F2**) | March 2019 | 16 | 16 | 4 | 4 | 16 | Ion Torrent sequencing (ITS2) |
| Corrèze (TMEL) | Forest 3 (**F3**) | May 2016 | 59 | 59 | 7 | 7 | - | Ion Torrent sequencing (ITS2) |
| Jardin des Plantes (JE) | Meadow 1 (**M1**) | June 2017 | 48 + 42 | 46 | 11 | 10 | - | Ion Torrent sequencing (ITS2) |
| Gaillac | Edge 1 (**MF1**) | June 2021 | 141 | 141 | 30 | 13 | - | Illumina sequencing (ITS2) |
| | | September 2021 | 59 | 21 | 7 | 6 | - | Illumina sequencing (ITS2) + *FISH*-microscopy |
| | | July 2023 | 4 root systems | 4 | 2 | 2 | - | *FISH*-microscopy |
| | Edge 2 (**MF2**) | June 2021 | 38 | 38 | 10 | 7 | - | Illumina sequencing (ITS2) |
| | | September 2021 | 50 | 17 | 7 | 6 | - | Illumina sequencing (ITS2) + *FISH*-microscopy |
| | Forest 4 (**F4**) | June 2021 | 16 | 16 | 4 | 4 | - | Illumina sequencing (ITS2) |
| | Forest 5 (**F5**) | | 13 | 13 | 7 | 4 | - | |
| | Meadow 2 (**M2**) | | 41 | 41 | 10 | 6 | - | |
| **Total** | | | **557** | **433** | **42** | **17** | **34** | |

**<u>Supplementary Table 2:</u> Plant host family and site of sampling influence both the total root mycobiota composition and the EcM mycobiota composition of non-EcM plants.** Outputs of the PERMANOVA analysis (*adonis2* function *vegan* R package), performed using the Bray-Curtis distances computed from both relative abundance and Hellinger transformed data of the total fungal community (**a**, **b**) or the EcM community (**c**,**d**) only with 10 000 permutations. The analyses were performed on the samples collected in **(a), (c)** the areas producing *Russula* spp. fruitbodies and the adjacent forest and meadow and in **(b), (d)** four natural environments across France. *family* corresponds to the plant host family. *environment* corresponds to the area where samples from (a), (c) were collected (areas producing *Russula* fruitbodies, forest or meadow). *location* corresponds the four natural environments of (b), (d) (cf. Table S1).

| | **Total fungal community** | | | | | | |
|---|---|---|---|---|---|---|---|
| **(a)** | **Variable** | **$R^2$** | **p-value** | **(b)** | **Variable** | **$R^2$** | **p-value** |
| Relative abundance | family | 0.15 | $10^{-3}$ (***) | Relative abundance | family | 0.32 | $10^{-4}$ (***) |
| | site | 0.06 | $10^{-3}$ (***) | | site | 0.05 | $10^{-4}$ (***) |
| | interaction | 0.06 | $10^{-4}$ (***) | | interaction | 0.01 | $6 \times 10^{-4}$ (***) |
| | Residuals | 0.73 | - | | Residuals | 0.63 | - |
| Hellinger transformation | family | 0.17 | $10^{-4}$ (***) | Hellinger transformation | family | 0.37 | $10^{-4}$ (***) |
| | site | 0.08 | $10^{-4}$ (***) | | site | 0.06 | $10^{-4}$ (***) |
| | interaction | 0.06 | $10^{-4}$ (***) | | interaction | 0.01 | $1.9 \times 10^{-3}$ (***) |
| | Residuals | 0.69 | - | | Residuals | 0.57 | - |
| | **EcM fungal community** | | | | | | |
| **(c)** | | **$R^2$** | **p-value** | **(d)** | | **$R^2$** | **p-value** |
| Relative abundance | family | 0.12 | $10^{-4}$ (***) | Relative abundance | family | 0.34 | $10^{-4}$ (***) |
| | site | 0.03 | $10^{-4}$ (***) | | site | 0.08 | $10^{-4}$ (***) |
| | interaction | 0.04 | $10^{-4}$ (***) | | interaction | 0.01 | 0.09 |
| | Residuals | 0.82 | - | | Residuals | 0.57 | - |
| Hellinger transformation | family | 0.13 | $10^{-4}$ (***) | Hellinger transformation | family | 0.36 | $10^{-4}$ (***) |
| | site | 0.04 | $10^{-4}$ (***) | | site | 0.09 | $10^{-4}$ (***) |
| | interaction | 0.04 | $10^{-4}$ (***) | | interaction | 0.01 | 0.05 |
| | Residuals | 0.78 | - | | Residuals | 0.57 | - |

**Supplementary Table 3: Bipartite networks metrics and significance.**

We constructed bipartite networks of interactions between EcM fungi and non-EcM plants at two taxonomic levels: fungal genera and plant families **(a)** fungal OTUs and plant species **(b)**. At each taxonomic level, we constructed one network for the Gaillac sites (local scale), one network for the sites across France (regional scale), and one network per site. We characterized their structure using several network metrics (specialization, modularity and weighted nestedness) computed with the *bipartite* R package. We then tested whether obtained values are significantly different than expected by chance using null models (Methods S3.3).

| (a) | Genera/Families | | | |
|---|---|---|---|---|
| | $H_2$’ | Q | wNODF | C |
| **France** | **0.80*** | **0.56*** | **14.6* (-)** | 0.27 |
| M1 | 0.89 | 0.35 | 1.35 | 0.67 |
| F1 | **0.87*** | **0.59*** | 36.7 | 0.63 |
| F2 | 0.59 | 0.41 | 45.5 | 0.52 |
| F3 | 0.88 | 0.48 | **32.2* (+)** | 0.39 |
| **Gaillac** | **0.44*** | **0.33*** | **47.5* (+)** | 0.43 |
| M2 | **0.53*** | **0.45*** | 51.6 | 0.60 |
| F4 | **0.50*** | **0.41*** | 44.5 | 0.68 |
| F5 | **0.85*** | **0.45*** | **30.9* (-)** | 0.63 |
| MF1 | **0.43*** | **0.27*** | **51.3* (+)** | 0.49 |
| MF2 | 0.38 | 0.26 | 40.6 | 0.47 |

| (b) | OTUs/Species | | | |
|---|---|---|---|---|
| | $H_2$’ | Q | wNODF | C |
| **France** | **0.85*** | **0.73*** | 2.94 | 0.10 |
| M1 | **0.82*** | **0.58*** | 19.9 | 0.37 |
| F1 | 0.65 | 0.43 | 5.12 | 0.48 |
| F2 | **0.77*** | **0.51*** | 5.64 | 0.47 |
| F3 | **0.82*** | 0.52 | 5.71 | 0.32 |
| **Gaillac** | **0.64*** | **0.55*** | **22.1* (+)** | 0.20 |
| M2 | **0.73*** | **0.59*** | 8.32 | 0.28 |
| F4 | **0.77*** | **0.52*** | 9.83 | 0.52 |
| F5 | **0.89*** | **0.72*** | 4.75 | 0.30 |
| MF1 | **0.71*** | **0.62*** | **19.7* (+)** | 0.25 |
| MF2 | **0.69*** | **0.58*** | **22.7* (+)** | 0.40 |

**<u>Supplementary Table 4:</u> Percentage of OTUs shared (or not) between rhizosphere and roots per species and their contribution to the EcM community.**

For each species, we determined the percentage of OTUs found only in rhizosphere samples, shared between rhizosphere and roots and found only in roots (number on the left). We also computed the proportion of the EcM community that these represent (number on the right). For instance, 26% of EcM OTUs are shared between rhizosphere and roots samples of *A. maculatum*, but these represent 66% of the total EcM reads.

| | Rhizosphere (%) | Shared (%) | Roots (%) | Total number of EcM OTUs |
|---|---|---|---|---|
| *Arum maculatum* | 73 / 34 | 26 / 66 | 1.3 / <1 | 74 |
| *Convallaria majalis* | 29 / 20 | 60 / 80 | 10 / <1 | 48 |
| *Euphorbia* sp. | 49 / 7 | 49 / 93 | 2.3 / <1 | 43 |
| *Ficaria verna* | 30 / 31 | 62 / 69 | 8 / <1 | 37 |
| *Galium aparine* | 51 / 8 | 38 / 81 | 11 / 11 | 37 |
| *Mercurialis perennis* | 47 / 15 | 50 / 85 | 2.8 / <1 | 36 |
| *Poa* sp. | 25 / <1 | 65 / 99 | 10 / <1 | 40 |

**<u>Supplementary Table 5:</u> Rhizosphere and roots have similar EcM mycobiota composition.**

Outputs of the PERMANOVA analysis (*adonis2* function *vegan* R package) performed using both relative abundance and Hellinger transformed data with 10 000 permutations. The analyses were performed on the samples collected in the ORS and ORY sites. (a) Analysis of the total fungal community. (b) Analysis of the EcM fungal community. *species* corresponds to the plant host species and *compartment* to the plant compartment (rhizosphere or root). *interaction* is the interaction term between the two variables (*species* x *compartment*).

| (a) | Variable | $R^2$ | p-value | (b) | Variable | $R^2$ | p-value |
|---|---|---|---|---|---|---|---|
| **Relative abundance** | species | 0.16 | $10^{-4}$ (***) | **Relative abundance** | species | 0.27 | $10^{-4}$ (***) |
| | compartment | 0.06 | $10^{-4}$ (***) | | compartment | 0.02 | 0.12 |
| | interaction | 0.12 | $10^{-4}$ (***) | | interaction | 0.06 | 0.88 |
| | Residuals | 0.66 | - | | Residuals | 0.65 | - |
| **Hellinger transformation** | species | 0.22 | $10^{-4}$ (***) | **Hellinger transformation** | species | 0.23 | $10^{-4}$ (***) |
| | compartment | 0.10 | $10^{-4}$ (***) | | compartment | 0.01 | 0.75 |
| | interaction | 0.10 | $10^{-4}$ (***) | | interaction | 0.06 | 0.98 |
| | Residuals | 0.57 | - | | Residuals | 0.70 | - |

**<u>Supplementary Table 6:</u> Results of the sensitivity and specificity test of the newly designed probes RUS899 and RUS101.**

In order to test for the sensitivity and specificity of the newly designed probes RUS101 and RUS899, we hybridized fungi collected from pure cultures or from samples collected *in situ*. + and – signs characterize the intensity of the fluorescence signal.

| **Tested probes** | ***Phanerochaete* sp.** | ***Lactarius quietus*** | ***Russula luteotacta*** | ***Russula* sp. *in vitro*** | ***Russula* sp. *in situ*** | ***Fungi* sp. 2 C2** | ***Fungi* sp. 1 C1** |
|---|---|---|---|---|---|---|---|
| | Pure cultivated hyphae | Pure cultivated hyphae | Pure cultivated hyphae | Pure cultivated hyphae | Harvested ascocarp (cape) | Harvested Ascocarp | Harvested Ascocarp |
| EUK516 | ++ | ++ | ++ | ++ | ++ | ++ | ++ |
| RUS101 | - | +/- | + | - | + | NA | NA |
| RUS899 | - | +(+) | + | +(+) | ++ | - | - |
| NONRUS506 | NA | NA | NA | NA | - | - | -/+ |

**<u>Supplementary Table 7:</u> Details on plant host investigated for *FISH* detection collected.**

Information regarding potential plant host investigated for the presence of *Russula* spp. hyphae. Plant species, sampling date, probes used for hybridization are given and whether *Russula* spp. hyphae could be identified within the roots.

| **Plant species** | **Sampling date** | **Probes** | **Detection of *Russula* spp. hyphae (Y/N)** |
|---|---|---|---|
| ***Ranunculus bulbosus*** | Sept. 2021 | Rus899-ATTO633/Euk51-ATTO565 & Rus101-ATTO633/Euk51-ATTO565 | Y |
| ***Carex* sp.** | Sept. 2021 | Rus899-ATTO633/Euk51-ATTO565 | N |
| ***E. amygdaloides*** | Sept. 2021 | Rus899-ATTO633/Euk51-ATTO565 | N |
| ***P. officinarium*** | Sept. 2021 | Rus899-ATTO633/Euk51-ATTO565 | N |
| | July 2023 | Rus101-ATTO633/Euk51-ATTO565 | Y/N |
| ***Carduus pycnocephalus*** | July 2023 | Rus101-ATTO633/Euk51-ATTO565 | Y |

### Supplementary Figures (1-11)

<u>**Supplementary Figure 1:**</u> **Location of the sampling sites and details of the Gaillac sites.** We sampled roots of non-EcM plant species in areas in several sites across France in forests (F) and meadows (M). In Gaillac, we also collected plants in edges of forests where abundant fruitbodies of *Russula* spp. were observed (MF). **(a)** Location of the sites in France. **(b)** Details on the location of the five Gaillac sites. Scale bar: 400 m. © IGN **(c)** Area producing *Russula* spp. fruitbodies (MF) and the adjacent forest (F) and meadow (M). **(d)** *Russula* sp. fruitbodies observed in June 2021 when sampling non-EcM roots. F: Forest; M: Meadow; MF: forest edge.

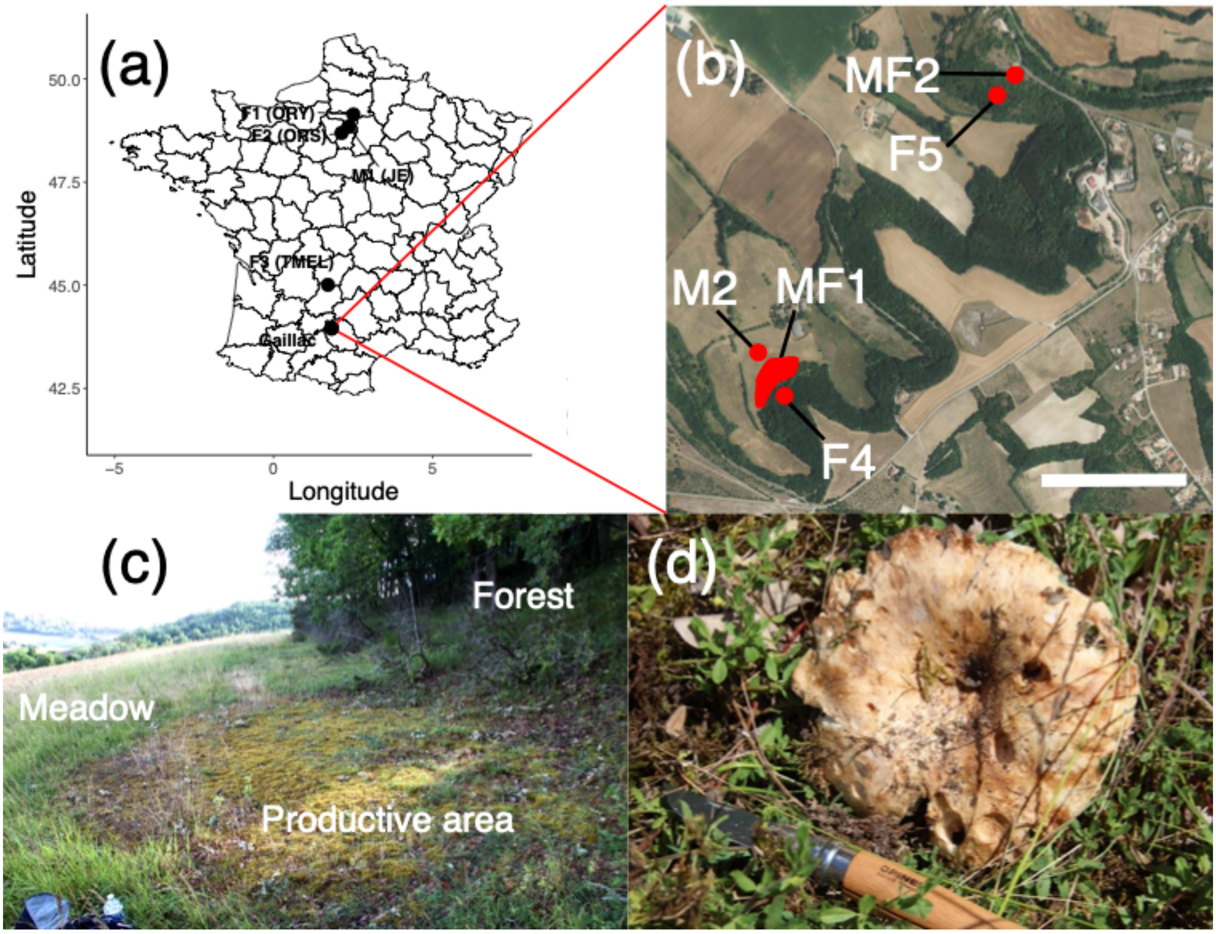

**<u>Supplementary Figure 2:</u> Total root mycobiota of non-EcM plants harvested in several sites in France.**

We described the root mycobiota of several non-EcM plants species in 9 sites across France using ITS2 amplicon sequencing. **(a)** Root mycobiota of plants collected in four sites across France. F: forest; M: meadow. **(b)** Root mycobiota of plants collected in five sites near Gaillac, France. MF: sites at the edge of the forest; F: forest; M: meadow.

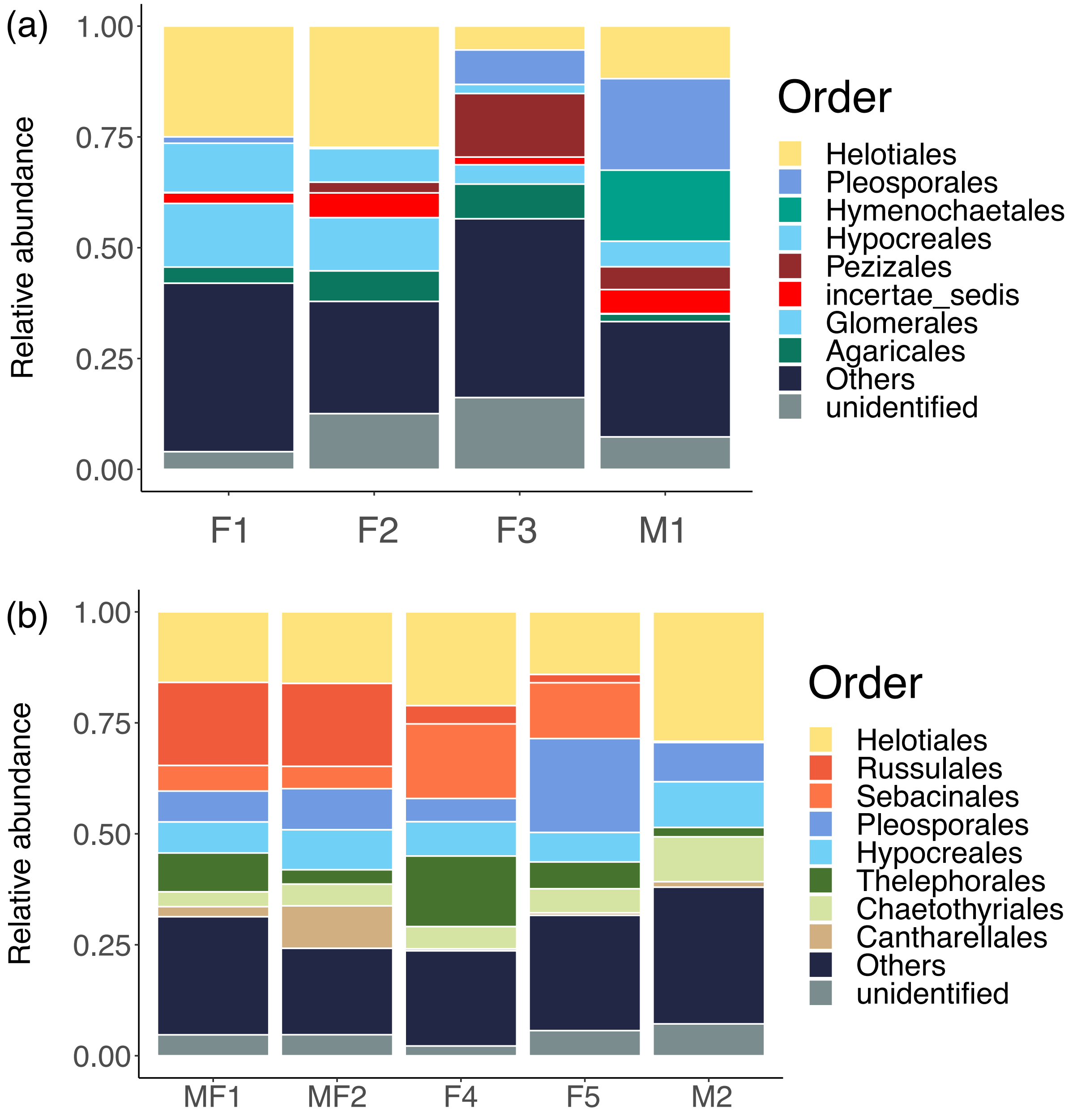

**<u>Supplementary Figure 3:</u> The proportion of EcM fungi in roots of non-EcM plant roots varies according to the plant host family and site.**

Boxplot of the proportion of EcM fungi by host family and by location. **(a)** Non-EcM plants collected in the four sites across France. **(b)** Plants collected in the five Gaillac sites.

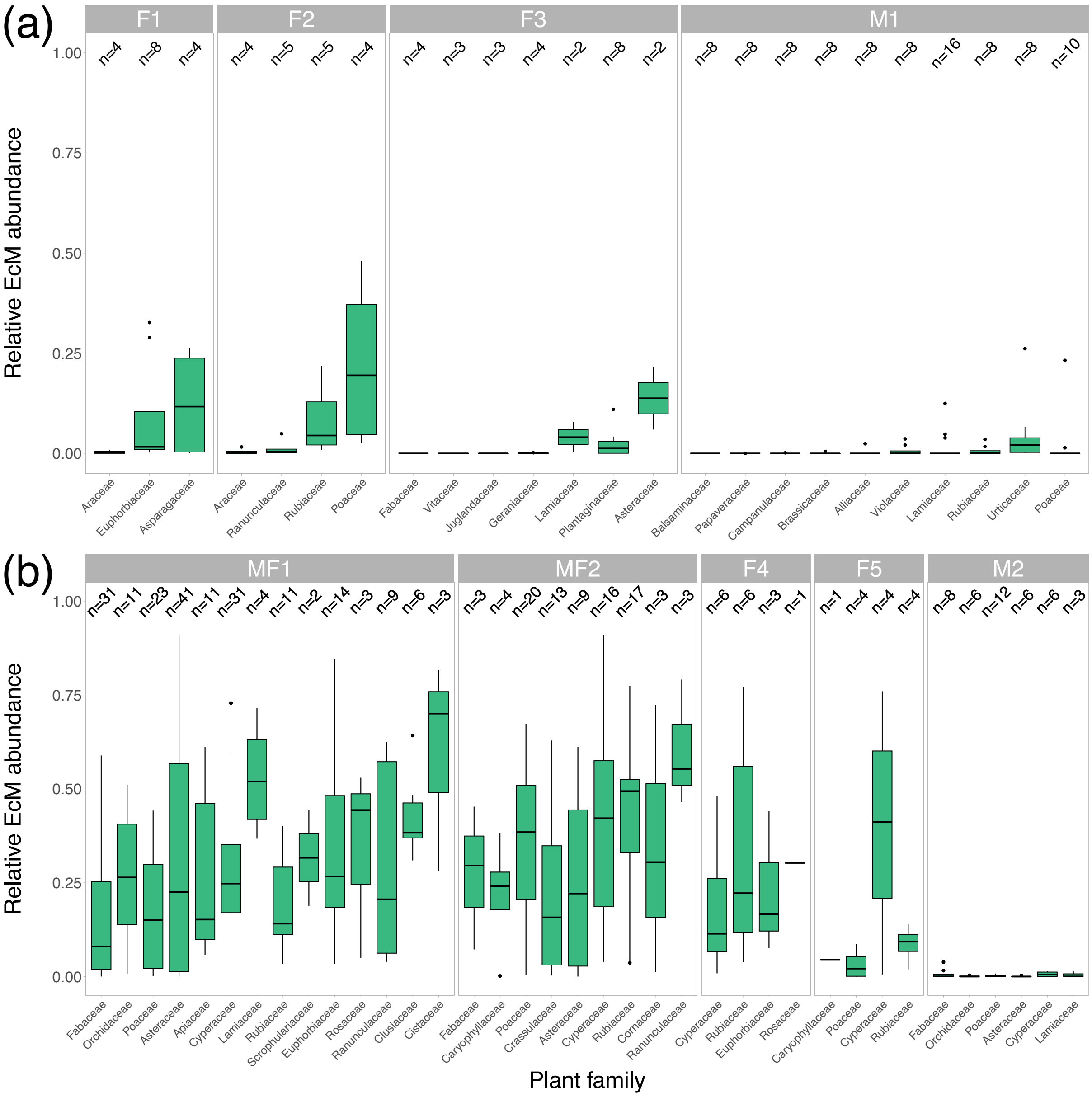

**<u>Supplementary Figure 4:</u> Forests, meadow and forest edges from Gaillac sites are characterized by OTUs differentially abundant.**

Differentially abundant OTUs between the three environment types of Gaillac sites: forest, meadow and forest edges. The x-axis corresponds to the Linear Discriminant Analysis score (LDA score) computed with the *LEfSE* method, and the size of the dots is proportional to the *p-value*: *-log(p-value)*.

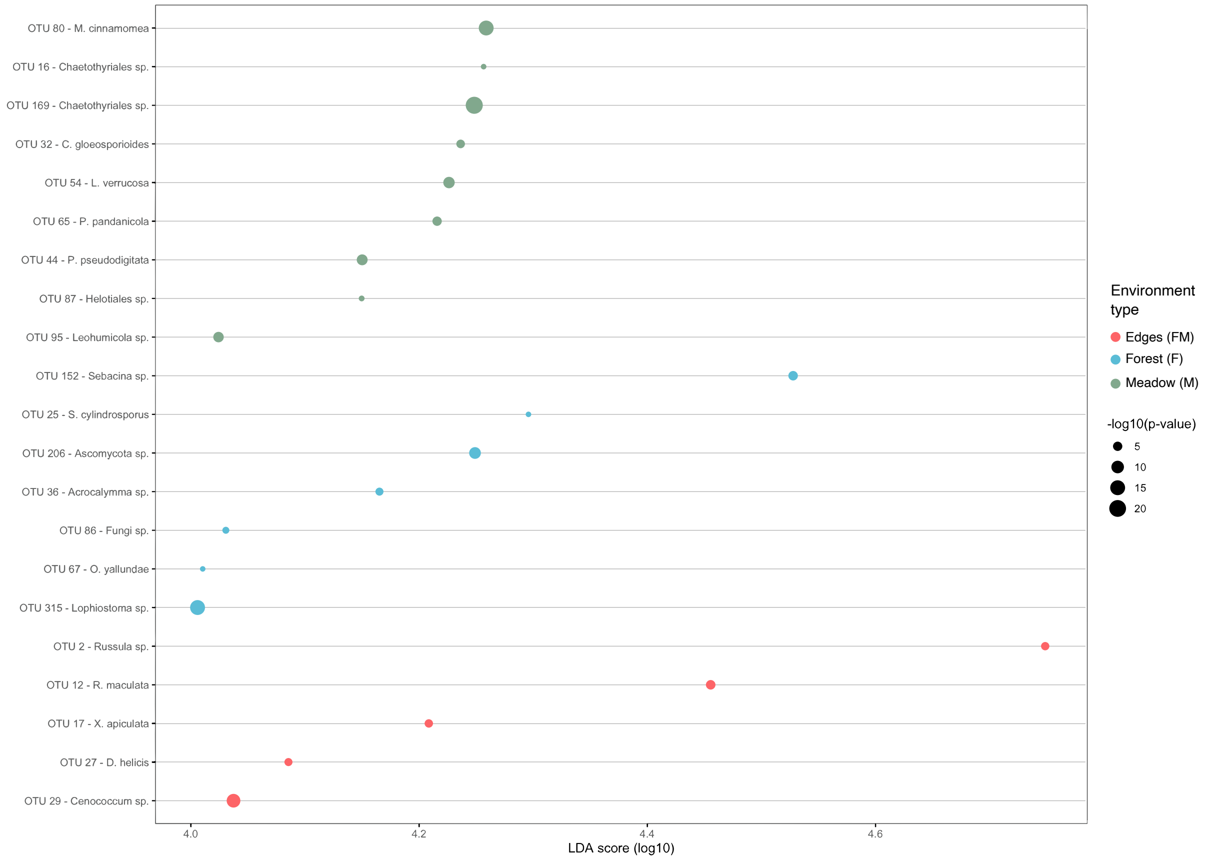

**<u>Supplementary Figure 5:</u> Bipartite network of interactions between plant families and fungal EcM genera across the four sites in France (F1, F2, F3 and M1).**

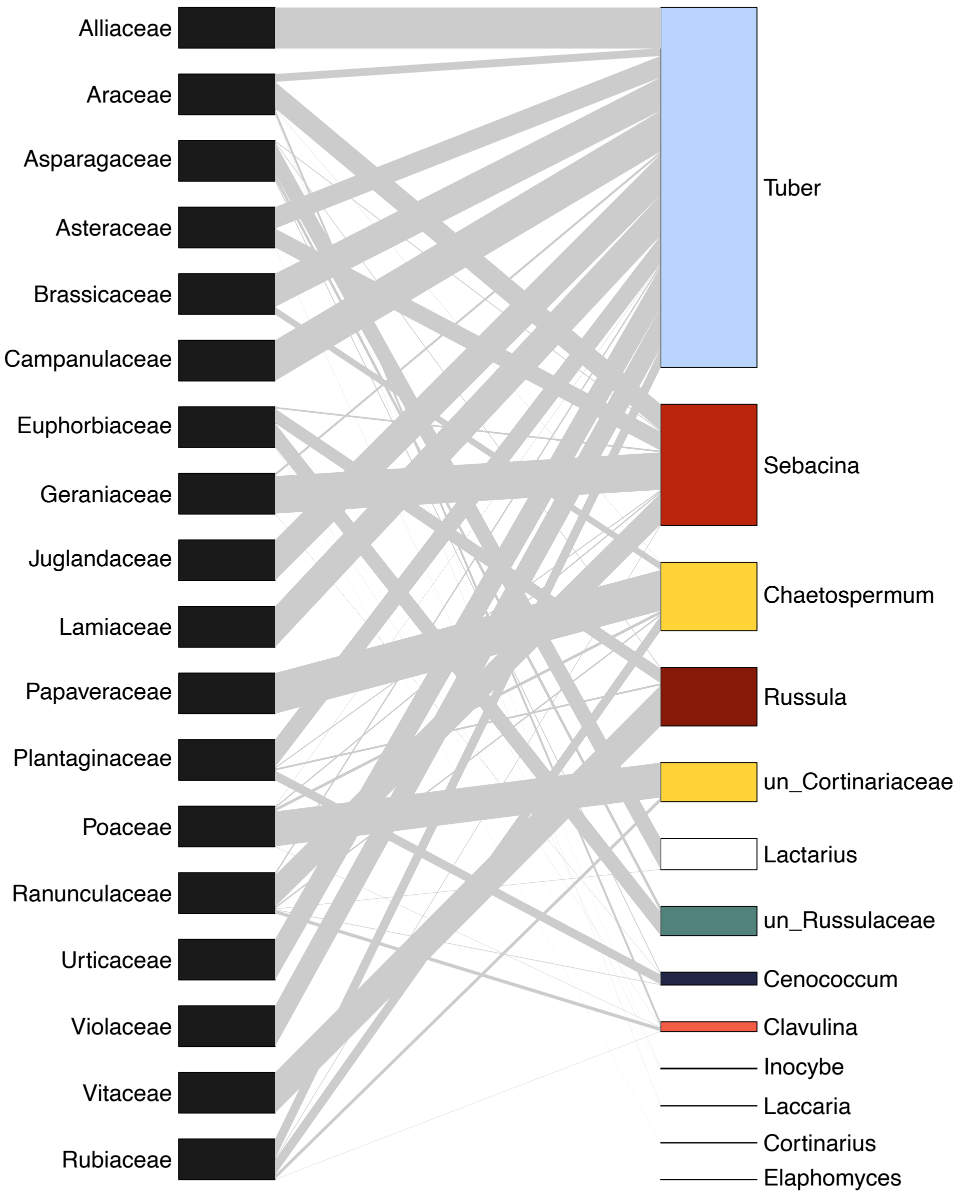

**<u>Supplementary Figure 6:</u> Root and rhizosphere have similar EcM community composition and diversity.**

**(a)** EcM community composition of the rhizosphere and the roots of species collected in the F1 and F2 sites. **(b)** Shannon index of rhizosphere and root EcM community for the seven species collected in F1 and F2 sites. Different letters indicate significant differences in EcM diversity (post-hoc Tukey's test).

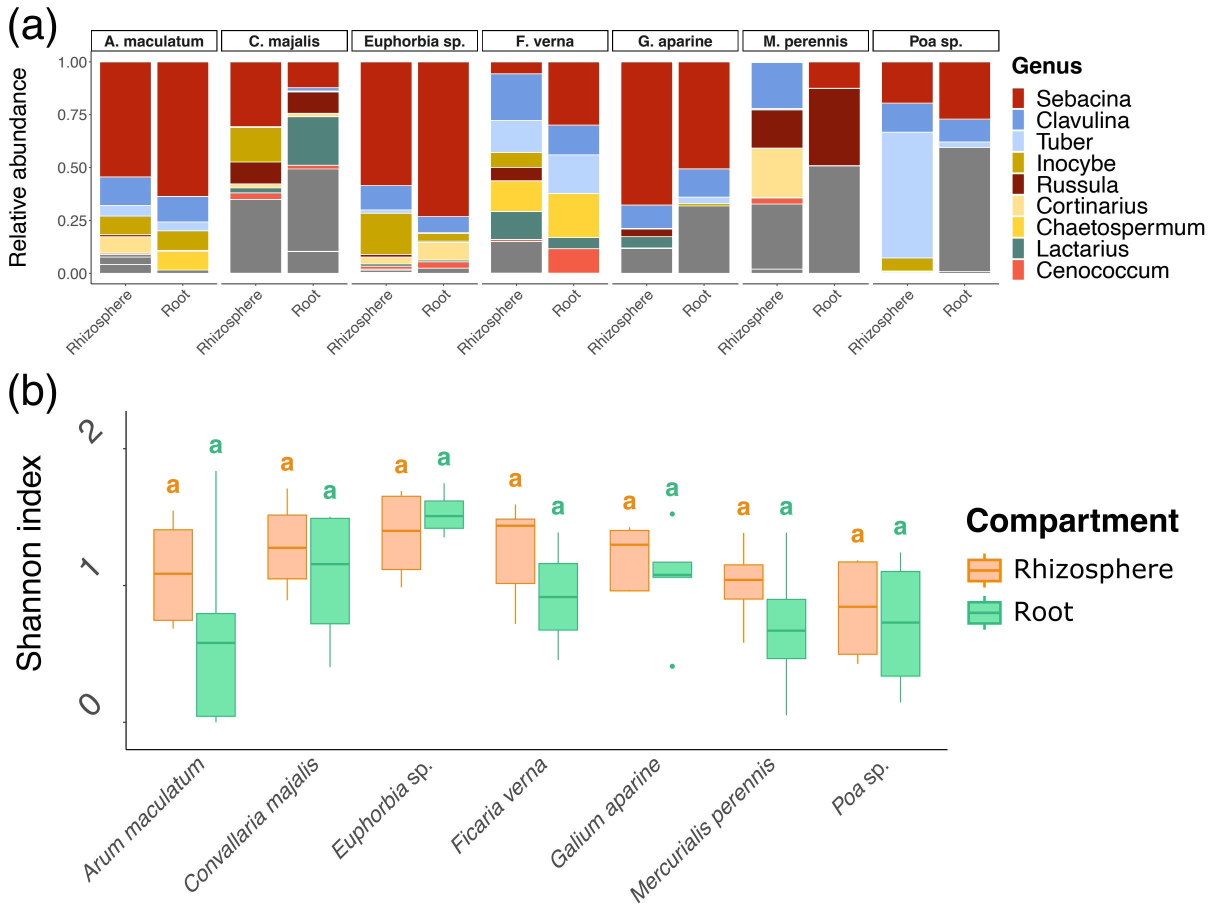

**<u>Supplementary Figure 7:</u> Some EcM genera are differentially abundant in roots and rhizosphere.** EcM genera significantly more abundant in roots or rhizosphere as computed using LEFsE procedure and their associated Linear Discriminant Analysis score (LDA; x-axis). The size of the dots corresponds to the log10 of the p-value computed by LEFsE.

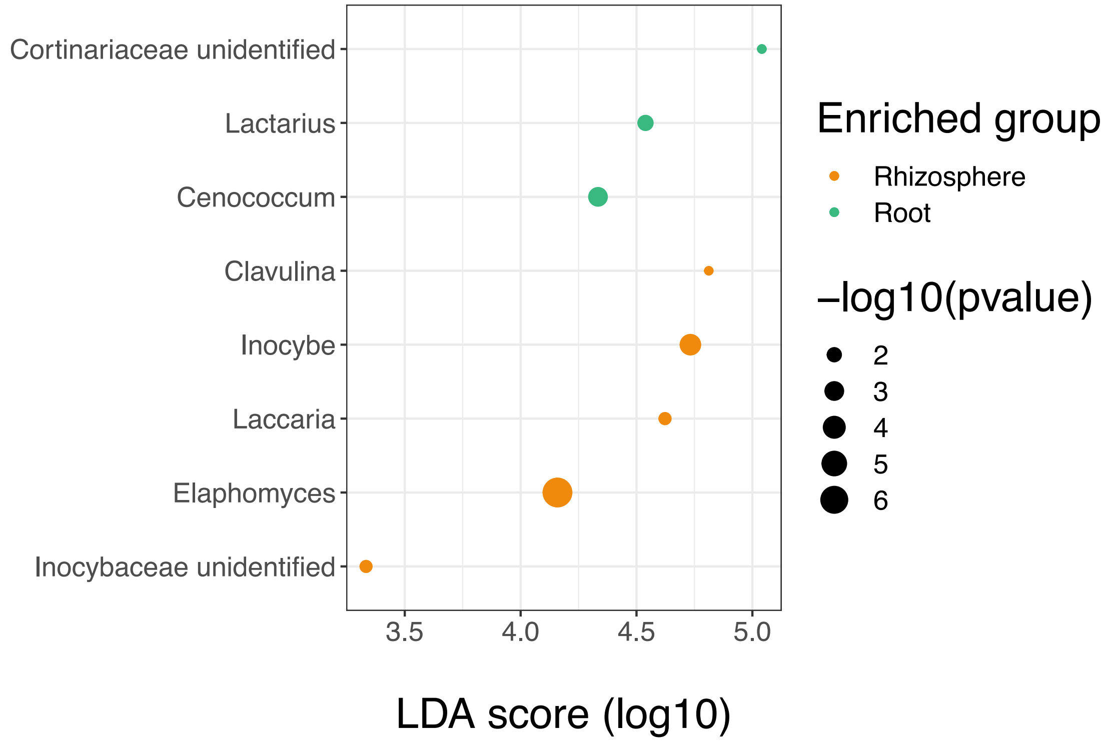

**<u>Supplementary Figure 8:</u> The newly designed probes Rus889 and Rus101 hybridize *in situ* with fungi from the genus *Russula*.**

Evaluation of *FISH* probes targeting *Russula* 18S ribosomal RNA with ascocarps or cultivated hyphae fixed from *Russula* sp. (left panel) or *Phanerochaete* sp. (right panel), respectively. Co-hybridization experiments with (1) a non-specific probe targeting eukaryotic cells (EUK516, second column of each panel) as a positive control with high signal intensity in the cytoplasm of fungal hyphae in combination with (2) *Russula* probes; RUS899 (**a-b** and **c-d**) or RUS101 (**e-f** and **g-h**). For each co-detection, 2D images obtained from each single channel are presented. Hybridization with *Russula* probes (RUS899 or RUS101) showed selective hybridization on *Russula* ascocarp with a good signal intensity in fungal cytoplasm comparable to the positive probe EUK516, but not on hyphae from *Phanerochaete* culture. Hybridization with a nonsense probe (NonRUS899, **i-j** and **k-l**) or with a hybridization buffer without any *FISH* probe (**m** and **n**) demonstrates the absence of any fluorescence signal in the cytoplasm of fungal hyphae. *Russula* sense and nonsense probes are coupled with ATTO633 dye and EUK516 is coupled with ATTO565 dye. Identical laser power and gain detector were used for each image. Scale bars: 10µm.

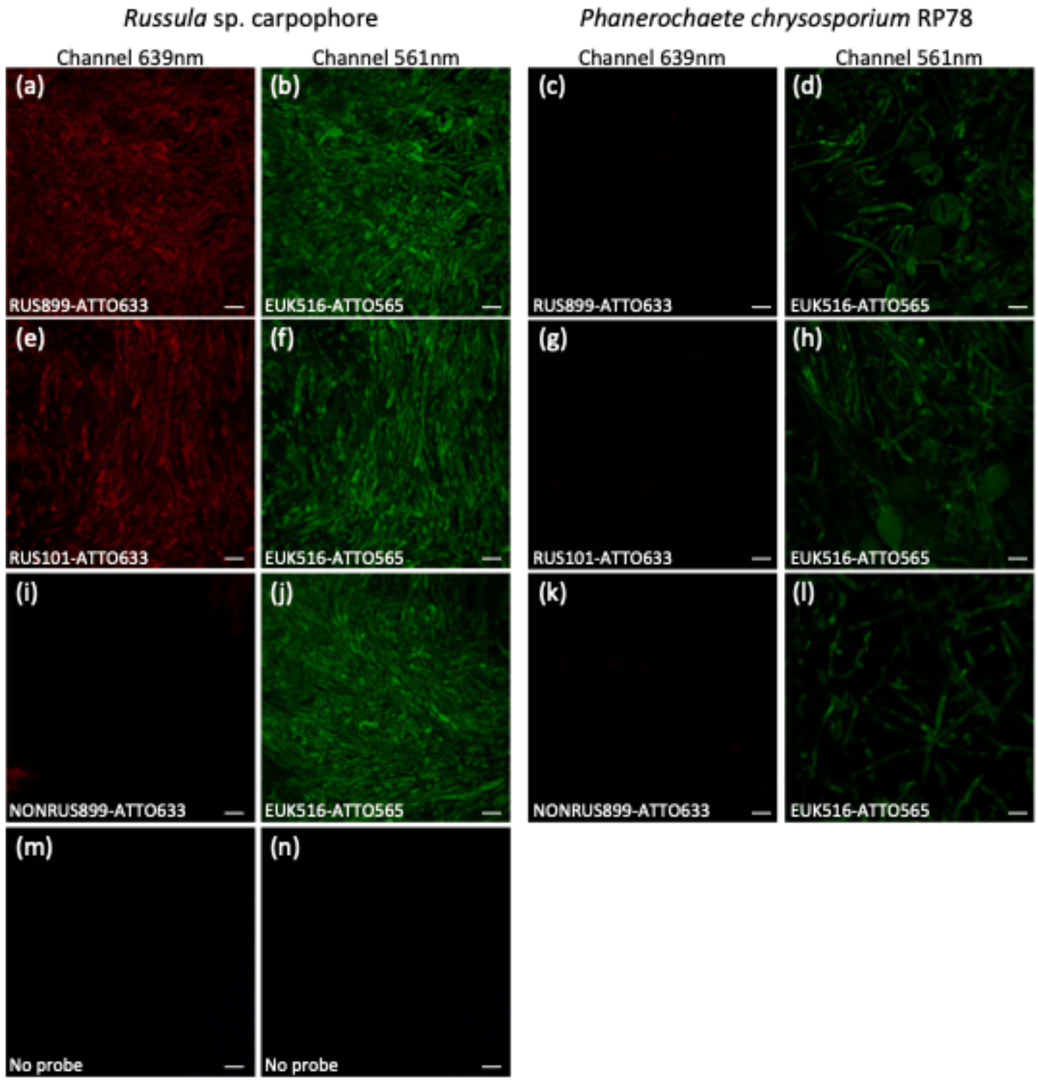

**<u>Supplementary Figure 9:</u> Observation of fungal hyphae in roots of *Carduus pycnocephalus* not hybridized with any *FISH* probe.**

**(a)**, **(b)** Negative controls to evaluate auto-fluorescence. Samples were processed with the same protocol of hybridization (Methods S4) but probes were replaced by sterile water (only the SR2000 fluorescent dye was kept). Most of the hyphae observed had no cytoplasmic autofluorescence in the ranges of the generalist probe EUK516 (561 nm, green signal) and the *Russula* probes RUS899/101 (639 nm, red signal), except in few cases where we could observe a fluorescent signal in the range of the EUK516 probe (green), as in (a). In (b), the red signal corresponds to plant tissues' autofluorescence at 639 nm. Scale bars: 30 µm.

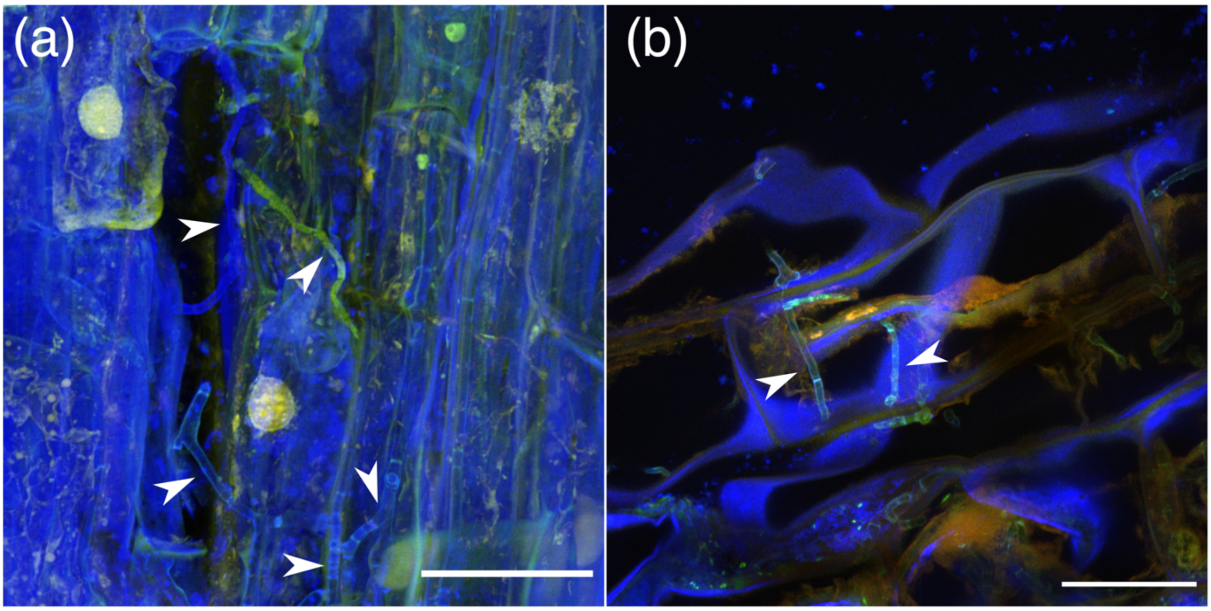

**<u>Supplementary Figure 10:</u> Observation of *Russula* spp. hyphae in the roots of *Carduus pycnocephalus* and of hyphae of other fungal species with the sense and nonsense probes (controls).**

**(a)**, **(c)** Roots of *C. pycnocephalus* harboring *Russula* sp. hyphae co-hybridized with the *Russula*-specific probe RUS101-ATTO633 (red signal) and with the generalist probe EUK516-ATTO565 (green signal). Co-hybridization appears in yellow/orange. **(a)** Co-hybridized hyphae in the apoplast of plant cells. **(c)** Fungal hyphae only hybridized with the generalist probe (green; top arrow) and *Russula* sp. co-hybridized hyphae (bottom arrow). **(b)**, **(d)** Nonsense controls: roots of *C. pycnocephalus* hybridized with the nonsense *Russula* probe NonRUS899-ATTO633 and the generalist probe EUK516-565. **(b)** We detected hyphae hybridized with the generalist probe (green signal, bottom arrow) and not hybridized at all (blue signal, top arrow). **(d)** Autofluorescence in the red spectrum was sometimes observed (green signal from the generalist probe + red autofluorescence; white arrow). Scale bars: 30 µm.

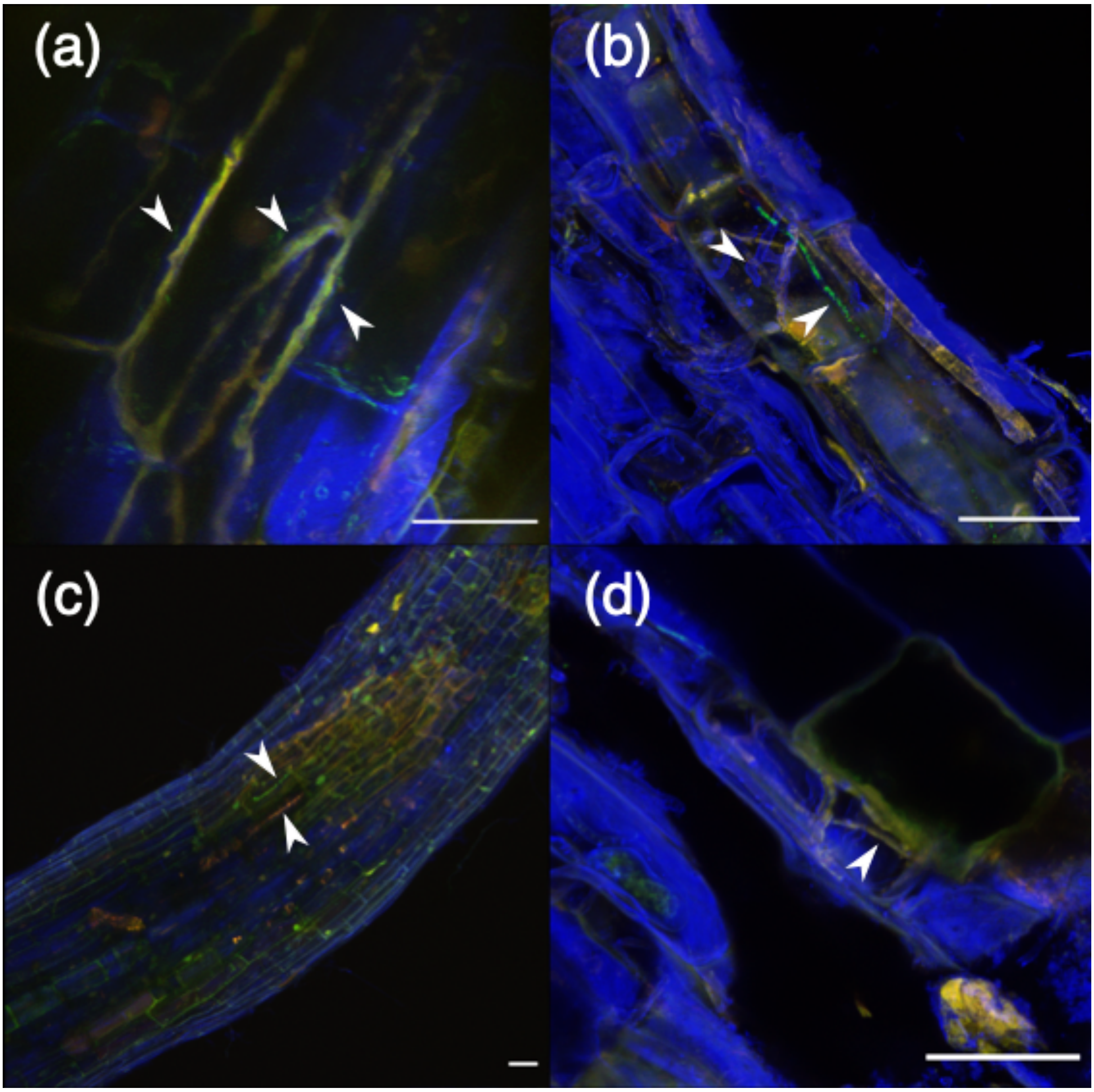

**<u>Supplementary Figure 11:</u> Observation of *Russula* spp. hyphae in the roots of Ranunculus bulbosus and of hyphae of other fungal species with the sense and nonsense probes (controls).** **(a)**, **(b)** Roots of R. bulbosus harboring *Russula* sp. hyphae hybridized with the *Russula*-specific probe RUS899-ATTO633 (red) and with the generalist probe EUK516-565 (green). Co-hybridization therefore appears yellow. **(c)** Roots of R. bulbosus harboring *Russula* sp. hyphae hybridized with the *Russula*-specific probe RUS101-ATTO633 (red) and with the generalist probe EUK516-565 (green). Co-hybridization therefore appears yellow. **(d)** Roots of R. bulbosus harboring a fungal hypha hybridized with the EUK516-565 (green) probe but not with the *Russula*-specific probe RUS101-ATTO633 (red). **(e)** Nonsense control: fungal hyphae in the roots of R. bulbosus hybridized with the nonsense *Russula* probe NonRUS899-ATTO633 (red signal) and the generalist probe EUK516-565 (green signal). Hyphae do not display any red signal (NonRUS899-ATTO633), showing that there is no unspecific binding from the probe. The red signal corresponds to plant cell wall autofluorescence. Scale bar: 30 µm.

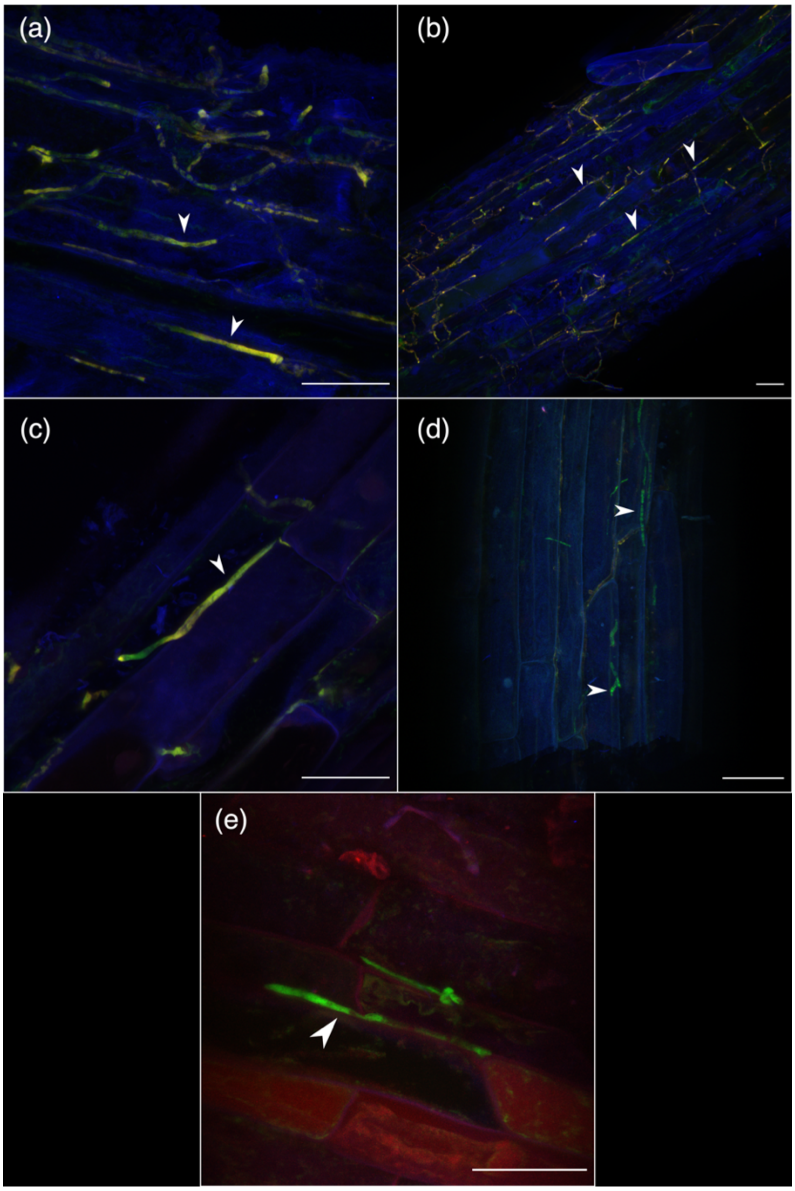
